# Molecular mechanism of attenuation of heat shock transcription factor 1 activity

**DOI:** 10.1101/803361

**Authors:** Szymon W. Kmiecik, Laura Le Breton, Matthias P. Mayer

**Author notes:** Corresponding author: Matthias P. Mayer, Center for Molecular Biology of Heidelberg University (ZMBH), DKFZ-ZMBH-Alliance, Im Neuenheimer Feld 282, 69120 Heidelberg, Germany.

## Abstract

The heat shock response is a universal transcriptional response to proteotoxic stress orchestrated by heat shock transcription factor Hsf1 in all eukaryotic cells. Despite over 40 years of intense research, the mechanism of HSF1 activity regulation remains poorly understood at a molecular level. In metazoa Hsf1 trimerizes upon heat shock through a leucin-zipper domain and binds to DNA. How Hsf1 is dislodged from DNA and monomerized remained enigmatic. Here, we demonstrate that trimeric Hsf1 is dissociated from DNA *in vitro* by Hsc70 and DnaJB1. Hsc70 acts at two distinct sites on Hsf1. Hsf1 trimers are monomerized by successive cycles of entropic pulling, unzipping the triple leucine-zipper. This process directly monitors the concentration of Hsc70 and DnaJB1. During heat shock adaptation Hsc70 first binds to the transactivation domain leading to partial attenuation of the response and subsequently, at higher concentrations, Hsc70 removes Hsf1 from DNA to restore the resting state.

## Introduction

The heat shock response (HSR) is an ancient transcriptional program, evolved in all organisms to cope with a wide variety of environmental, physiological and developmental stressful conditions that induce an imbalance of protein homeostasis. This transcriptional program up-regulates hundreds and down-regulates thousands of genes in metazoa ^1^. As the HSR was viewed as the paradigm for a homeostatic response, the mechanism of its activation and attenuation was intensively studied in the last four decades but still remains poorly understood at a molecular level ^2, 3^. The central role in regulating the HSR and restoring protein homeostasis in eukaryotic cells is fulfilled by the heat shock transcription factor 1 (Hsf1). Through this function Hsf1 is in metazoa at center stage of many physiological and pathophysiological processes like post-embryonic development and aging, cancer and neurodegeneration ^4–8^. Metazoan Hsf1 consists of a DNA binding domain (DBD), leucine zipper-like heptad-repeat regions A and B (HR-A/B) functioning as trimerization domain, a regulatory domain (RD), a third heptad repeat region (HR-C) and a transactivation domain (TAD) (Fig. 1A) ^9^. Recent structural studies in our laboratory revealed that RD and TAD are mostly unstructured ^10^.

**Figure 1:**
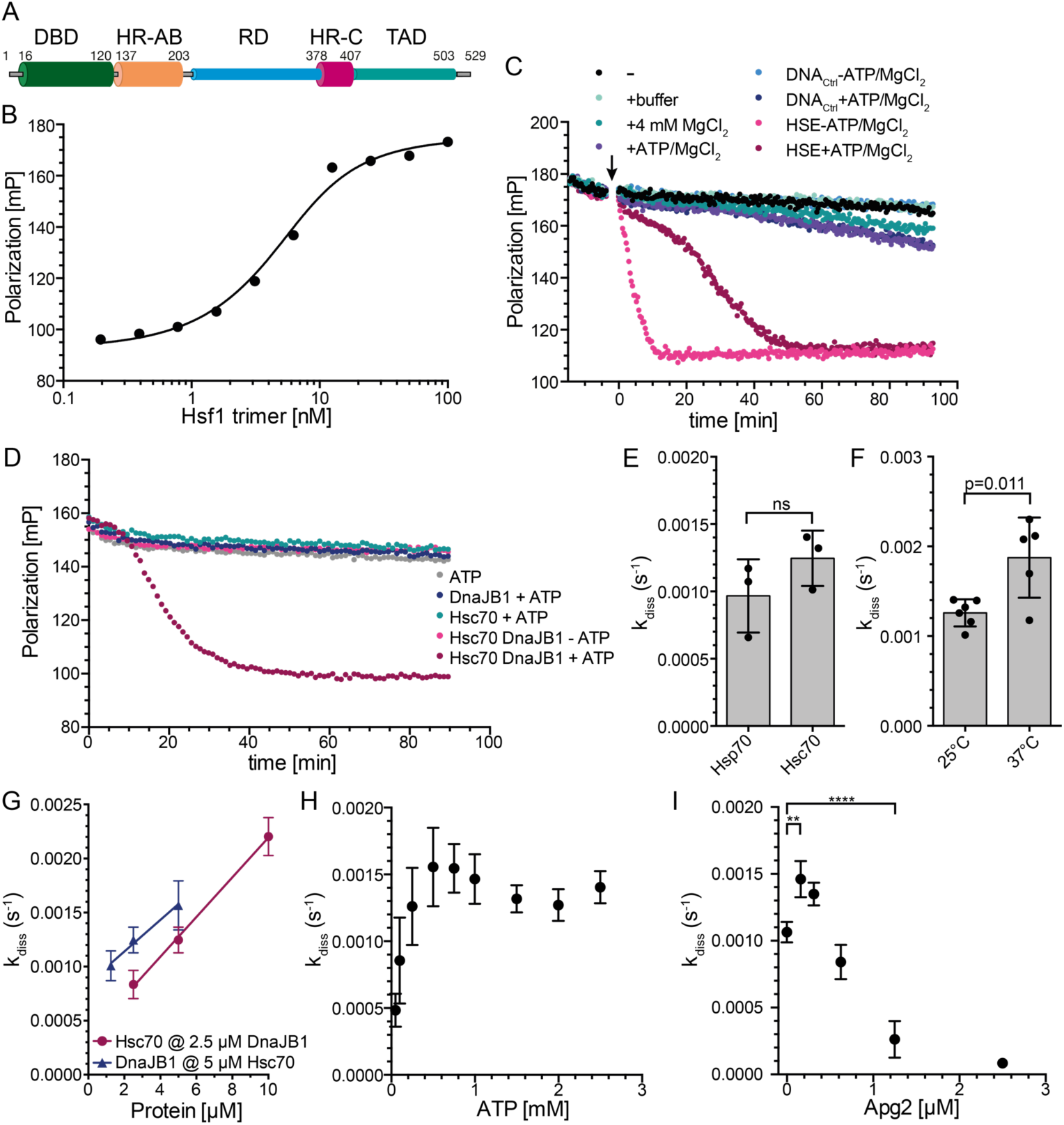
Hsf1 binds with high affinity but highly dynamic to heat shock elements (HSEs) and is dissociated from DNA by Hsc70 and DnaJB1. **A**, Domain structure of mammalian Hsf1. Intrinsically disordered regions are indicated with cylinders of smaller diameter. DBD, DNA binding domain, HR-AB; heptad repeat regions A and B; RD, regulatory domain, HR-C, heptad repeat region C; TAD, transactivation domain. **B**, Equilibrium titration of Hsf1 binding to Alexa 488-labeled HSE-DNA. Fluorescence polarization plotted versus the Hsf1 trimer concentration. K_D_ = 4.7 ± 1.9 nM (n = 4). **C**, Hsf1 rapidly switches between different HSE-containing double stranded DNA oligonucleotides. Trimeric Hsf1 was pre-incubated with Alexa 488-labeled HSE-DNA in MgCl_2_-free buffer. At timepoint 0 (arrow) buffer, 4 mM MgCl_2_, 2 mM ATP+4 mM MgCl_2_, control DNA (DNA_Ctrl_ at 10-fold molar excess over HSE-DNA), and HSE-containing DNA was added as indicated and fluorescence polarization followed over time. Shown is one of three identical experiments. **D**, Trimeric Hsf1 were bound to HSE-DNA and the indicated components added at timepoint 0. A representative experiment is shown. **E**, Hsp70 dissociates Hsf1 from DNA with similar rates as Hsc70. Rates were determined as shown in Fig. S1C. **F-I**, The rate of Hsc70/DnaJB1-mediated dissociation of Hsf1 from HSE-DNA depends on temperature (F), concentration of Hsc70 and DnaJB1 (G), concentration of ATP (H), and concentration of the nucleotide exchange factor Apg2 (HSPA4) (I).

In non-stressed mammalian cells Hsf1 is in monomer-dimer equilibrium in the nucleus and the cytosol ^10, 11^. During stress conditions Hsf1 accumulates in the nucleus and forms oligomers that gain increased affinity for binding to so-called heat shock elements (HSE: nGAAn) arranged in inverted repeats of three and more units ^12–14^. Hsf1 HR-C region serves as an intramolecular Hsf1 oligomerization repressor that operates as a temperature sensor, integrating over temperature and time at elevated temperatures by local unfolding and dissociation from HR-A/B ^9, 10^. The setpoint of this temperature rheostat is dependent on the concentration of Hsf1, allowing nuclear transport and Hsf1 expression to alter the fraction of trimerized Hsf1 at any given temperature ^10, 15–17^.

Based on accumulated knowledge several Hsf1 activation/attenuation mechanisms have been proposed ^18^. The most prominent Hsf1 activity regulation model, the chaperone titration model, assumes that Hsf1 is sequestered and inactivated by molecular chaperones; Hsf1 activation follows the recruitment of the chaperones to stress-denatured proteins. Since Hsp70 was found to co-precipitate with Hsf1, Hsp70 was implicated in repressing Hsf1 in the resting state or during attenuation ^19^. Other evidence suggested that Hsp70 is insufficient in metazoa for Hsf1 repression, contesting this model ^20^. Since loss of Hsp90 functionality activates the HSR, it was also suggested that Hsp90 chaperones sequester Hsf1 and keep it inactive ^21^. However, to demonstrate Hsp90 and Hsf1 interaction, crosslinking is required ^22^, unless an ATPase-deficient Hsp90 variant is used, indicating that Hsf1 interacts with Hsp90 only in the ATP-bound closed conformation ^23, 24^. In addition, *in vitro* and in the absence of cochaperones, Hsp90 favors Hsf1 trimerization and DNA binding ^10^. Although genetic evidence suggested an involvement of Hsp90 in HSR regulation also in yeast ^25^, more recent *ex vivo* data suggest that Hsp70 is associated with Hsf1 under non-stress conditions and this interaction is disrupted upon heat shock ^26, 27^. Whether such a model can be adopted for mammalian cells is not clear since Hsf1 is constitutively trimeric in yeast and does not rely on a monomer-trimer transition for activation ^28^. Moreover, the overall degree of sequence identity between yeast and human Hsf1 is just 17% and the proposed binding sites of Hsp70 in yeast Hsf1 are not conserved in human Hsf1.

Several different models have been proposed for HSR attenuation, Hsf1 dissociation from DNA and recycling of Hsf1. Hsp70 and its co-chaperone DnaJB1 have been suggested to bind to Hsf1 within the TAD (aa 425-439) thus attenuating Hsf1 activity by repressing the recruitment of the transcriptional machinery ^29^. Hsf1 acetylation in the DBD was proposed to be required for removing Hsf1 from DNA ^30^. Thereby the SIRT1 deacetylase plays a main role in delaying the attenuation of the HSR. The interaction between DNA and Hsf1 DBD relies on electrostatic contacts ^31^ and replacement of Lys80 and/or Lys118 in Hsf1 to glutamate significantly reduced DNA binding ^30^. Since the inhibition of proteasomal function also delayed HSR attenuation, it was suggested that, after acetylation-dependent dissociation, Hsf1 trimers are directed to proteasomal degradation ^32^. However, inhibition of the proteasome also leads to accumulation of misfolded proteins in the cytosol, eliciting the HSR. For *Drosophila* Hsf1 it was shown that trimers disassemble spontaneously to monomers at low concentrations ^33^. However, such a spontaneous dissociation was not observed for human Hsf1 ^10^.

In this work we demonstrate using purified components that both Hsc70 and Hsp70 in cooperation with DnaJB1 dissociate trimeric Hsf1 from DNA in the absence of Hsf1 acetylation. Furthermore, we show that during dissociation Hsf1 is monomerized, with most of Hsf1 remaining sequestered in complex with Hsc70. We identify several binding sites for Hsc70 within Hsf1, one of which in the transactivation domain involved in initial attenuation, a second close to the trimerization domain essential for Hsc70-mediated monomerization. We provide evidence that Hsc70-mediated monomerization of Hsf1 trimers occurs through stepwise unzipping of the triple leucine-zipper of the Hsf1-trimer by entropic pulling. Mutational alteration of the Hsc70 binding sites potentiates expression of a heat shock reporter in HSF1^-/-^ mouse embryonic fibroblasts. Based on these and published data we propose a comprehensive model for a dynamic regulation of Hsf1 activity that closely monitors availability of cellular Hsc70 and Hsp70.

## Results

### Hsf1 can migrate rapidly between different HSE-containing DNAs

To investigate DNA binding of purified human trimeric Hsf1, we used the previously established fluorescence polarization assay with Alexa488-labeled double stranded DNA oligonucleotides containing 3 inverted HSEs. We first titrated Hsf1 and established the dissociation equilibrium constant K_D_ to ≤ 5 nM (Fig. 1B) consistent with previous results ^10^. The interaction of Hsf1 with HSE-containing DNA was very stable and little decrease in fluorescence polarization was observed over 90 min (Fig. 1C) when only buffer, or excess unlabeled control DNA without HSEs was added. Addition of MgCl_2_ (4 mM) without or with ATP or excess of control DNA led to a slow decrease of polarization, presumably due to slow dissociation and aggregation of Hsf1 during the long incubation time. In contrast, if excess of unlabeled HSE-DNA was added in the absence of MgCl_2_ polarization decreased rapidly, reaching the polarization value of unbound labeled DNA within 10 min, indicating that Hsf1 can switch from one HSE-containing dsDNA segment to another at high rates. The decrease was significantly slower when the unlabeled HSE-DNA was added in the presence of MgCl_2_, suggesting the Mg^2+^ ions bound to the DNA backbone phosphate reduces the association rate of Hsf1 DBD to the DNA. From these experiments we conclude that individual DBDs of the Hsf1 trimer dissociate from the HSE-DNA and re-associate rapidly in the absence of Mg^2+^ ions and somewhat more slowly in the presence of Mg^2+^ ions. For further experiments we always pre-incubated trimeric Hsf1 with labeled HSE-containing dsDNA in the absence of MgCl_2_ to generate stably bound Hsf1-DNA complexes and then added different combinations of chaperones in the absence or presence of Mg^2+^·ATP.

### Hsc70 but not Hsp90 can dissociate Hsf1 from DNA

A recent publication proposed an inhibitory role of Hsp90 during the attenuation phase of the HSR ^24^. However, in our *in-vitro*-polarization DNA-binding assay neither human Hsp90α wild type nor its ATPase-deficient variant Hsp90α-E47A, which was shown to bind Hsf1 with higher affinity, had any influence on the change in polarization as compared to the controls (Fig. S1A), indicating that its effect during attenuation phase of the HSR was not achieved through dissociation of Hsf1 from DNA. This result is consistent with earlier findings that Hsp90 promotes Hsf1 trimerization and DNA binding ^10^.

In contrast, human Hsc70 in the presence of ATP and its J-domain cochaperone DnaJB1/Hdj1, which targets Hsc70 to client proteins by stimulating Hsc70’s ATPase activity, efficiently dissociated Hsf1 from DNA (Fig. 1D). This effect was not observed when any of the three components, Hsc70, DnaJB1 or ATP, was missing, or when Hsc70 wild type was replaced by its ATPase-deficient variant Hsc70-K71M, or its polypeptide binding defective variant Hsc70-V438F, or when DnaJB1 wild type was replaced by a variant (DnaJB1-H32Q,D34N) that is not able to stimulate Hsc70’s ATPase activity (Fig. S1B).

When analyzing the shape of the dissociation curve, we observed a short 5 to 10 min delay, during which the dissociation did not follow an exponential function, before the actual exponential dissociation phase started. The data were therefore fitted by a composite function and the rate only represents the exponential phase of the dissociation reaction (Fig. S1C). The dissociation rate was not significantly different whether we used Hsf1 purified from *E. coli* as trimer and not heat shocked or as monomer and subsequently heat shocked at 42°C for 10 min (Fig. S1D). Also, the heat inducible Hsp70 (HSPA1A/B) dissociated Hsf1 from DNA with similar rates as the constitutive Hsc70 (HSPA8) (Fig. 1E). The reaction was, as expected, temperature dependent and increasing the temperature from 25 to 37°C increased the dissociation rate significantly (Fig. 1F). The kinetics of Hsc70-mediated Hsf1 dissociation from DNA were very similar to the kinetics with which Hsf1-mediated transcription activation and DNA binding of Hsf1 decreased in HeLa cells during recovery after a heat shock ^34^. The dissociation reaction rate was strongly dependent on the concentration of Hsc70 and DnaJB1, increasing almost threefold between 2.5 and 10 µM Hsc70 at 2.5 µM DnaJB1 and 1.5-fold between 1.25 to 5 µM DnaJB1 at 5 µM Hsc70 (Fig. 1G).

The dissociation reaction was also strongly dependent on the ATP concentration between 0.05 to 0.5 mM, but not between 0.5 to 2.5 mM (Fig. 1H). In the presence of physiological concentrations of ATP, the life-time of an Hsc70-client protein complex is limited by nucleotide exchange that is accelerated by nucleotide exchange factors^35^. We therefore added the nucleotide exchange factor Apg2 to the dissociation reaction. At very low concentrations, Apg2 accelerated the dissociation reaction, but at higher concentrations it strongly inhibited the reaction and prevented Hsc70-mediated dissociation of Hsf1 from DNA (Fig. 1I). This is similar as Apg2 action in Hsc70-mediated protein disaggregation, where also low concentration of Apg2 stimulate and high concentrations inhibit the reaction ^36^. Taken together, Hsc70 and Hsp70 dissociate Hsf1 from its binding sites in promoter DNA in the presence of DnaJB1 and physiological concentrations of ATP in a strongly concentration dependent manner. This reaction is independent of Hsf1 acetylation in the DBD.

### Hsc70 dissociate Hsf1 from DNA by monomerization of Hsf1 trimers

For the Hsc70-mediated dissociation of Hsf1 from DNA different mode of actions are imaginable. In analogy to Hsp70 action on p53 ^37, 38^, Hsc70 could directly interact with the DBD of Hsf1 to competitively or allosterically remove the DBD from the DNA. Alternatively, Hsc70 could monomerize Hsf1, which would lead to dissociation from DNA because individual DBDs have only a very low affinity for the HSEs and the high affinity of Hsf1 for heat shock promoters is an avidity effect of three DBDs binding simultaneously. To test the second hypothesis, we followed the dissociation reaction by fluorescence polarization and took samples every 10 min to analyze the oligomeric state of Hsf1 by blue-native polyacrylamide gel electrophoresis (BN-PAGE) and immunoblotting with Hsf1 specific antisera (Fig. 2). In the beginning of the reaction only trimeric and higher order oligomeric Hsf1 was present. In the course of the dissociation reaction the trimer band disappeared and some Hsf1 monomer became visible. Most of Hsf1 exhibited an electrophoretic mobility that is in between the trimeric and the monomeric state, presumably bound to Hsc70. If any of the components were missing, the Hsf1 remained oligomeric. Quantification of the Hsf1 oligomer band and the rest of the Hsf1 species revealed that the oligomer band disappeared within experimental error with the same rate as Hsf1 dissociated from DNA in the fluorescence polarization assay, strongly arguing that Hsc70 dissociates Hsf1 from DNA by monomerization. Hsc70 also monomerized trimeric Hsf1 with the same rate in the absence of DNA (Fig. 2C).

**Figure 2:**
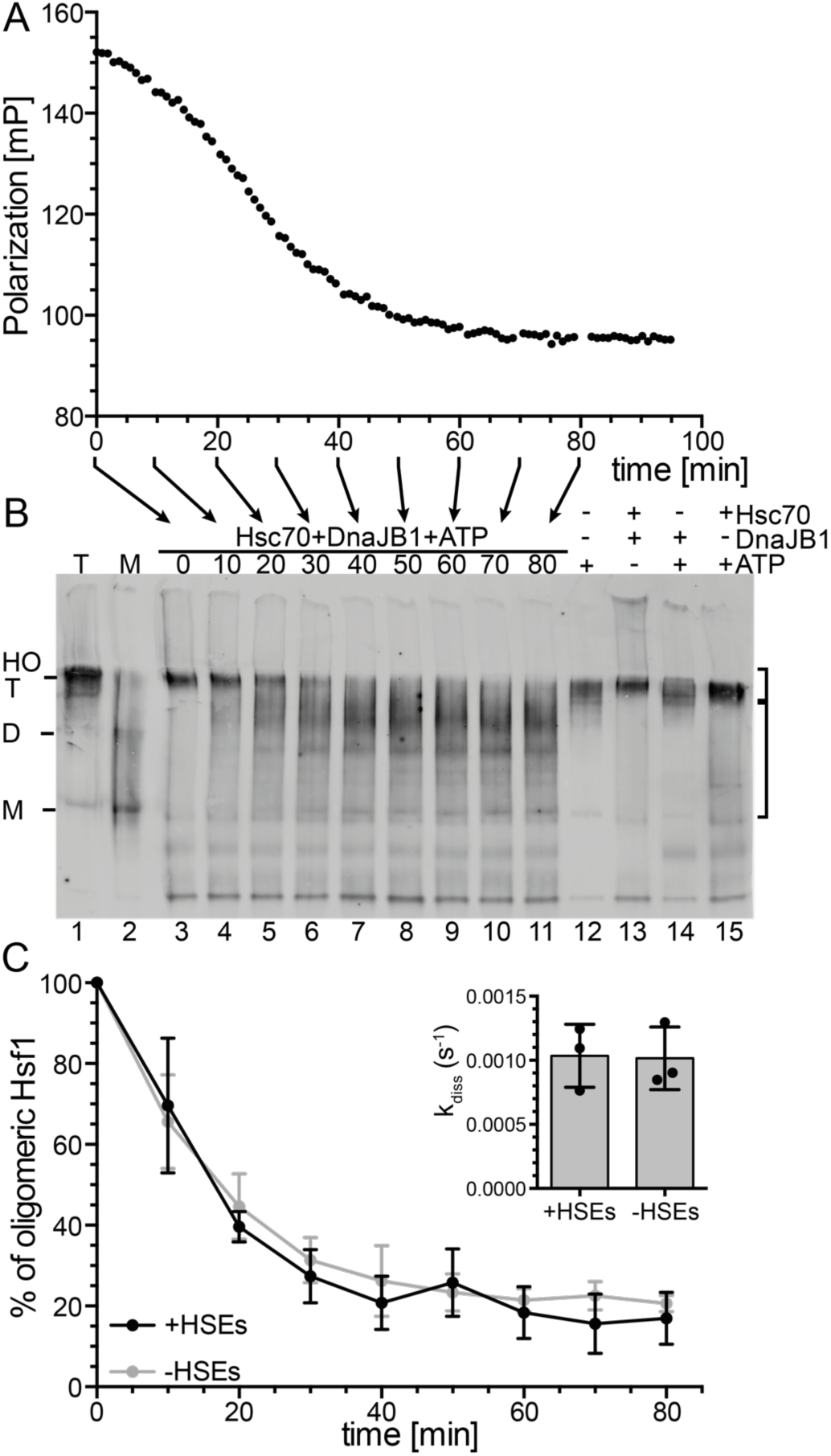
Hsc70 and DnaJB1 dissociate Hsf1 from HSE-DNA by monomerization of the Hsf1 trimer. **A-B**, Standard Hsc70/DnaJB1-mediated dissociation reaction of Hsf1 from Alexa488-labeled HSE-DNA monitored by fluorescence polarization (A). At the timepoints indicated by the arrows samples were taken and separated by blue-native polyacrylamide gel electrophoresis, blotted onto a PVDF membrane and detected with an Hsf1-specific antiserum (B). Lanes 1, purified Hsf1 trimer (T); 2, purified Hsf1 monomer (M); 3-11, samples from the Hsc70/DnaJB1-mediated Hsf1 dissociation reaction (0 to 80 min); 12-15, Dissociation reaction incubated for 80 min missing individual components as indicated. HO, higher order oligomers; T, trimer; D, dimer; M, monomer. **C**, Quantification of the amounts of Hsf1 species of the blot shown in B and two similar blots as indicated by the brackets to the right; upper bracket, DNA bound timers and higher order oligomers; lower bracket, monomers and Hsc70-bound species. Shown are means ± SD (n = 3).

### Hsc70 binds to several sites in monomeric and trimeric Hsf1 and destabilizes the trimerization domain

To elucidate how Hsc70 dissociates Hsf1 trimers, we first wanted to identify the binding site of Hsc70 within Hsf1 using hydrogen exchange mass spectrometry (HX-MS), which is suitable to detect solvent accessibility of amide hydrogens of the peptide backbone and thus conformational changes in proteins and protein-protein interactions ^39^, as described in detail previously ^10, 40^. Briefly, we pre-incubated Hsf1 in the absence or presence of Hsc70 or DnaJB1 or both for 0 or 30 min at 25°C, diluted the sample 1:10 in D_2_O containing buffer and incubated for 30 or 300 s at 25°C. Subsequently, the exchange reaction was quenched and the samples analyzed by LC-MS, including online digestion of the proteins by immobilized pepsin to localize the incorporated deuterons to specific segments of the protein.

Plotting the deuteron incorporation into Hsf1 in the presence of Hsc70 or DnaJB1 minus the deuteron incorporation in the absence of chaperones revealed 5 regions of significant protection that were observed for trimeric as well as for monomeric Hsf1, suggesting 5 potential binding sites (Fig. 3A, Fig. S2). Surprisingly, we did not detect any protection close to amino acids 395-439 previously proposed to harbor the Hsc70 binding site in Hsf1 ^29^.

**Figure 3:**
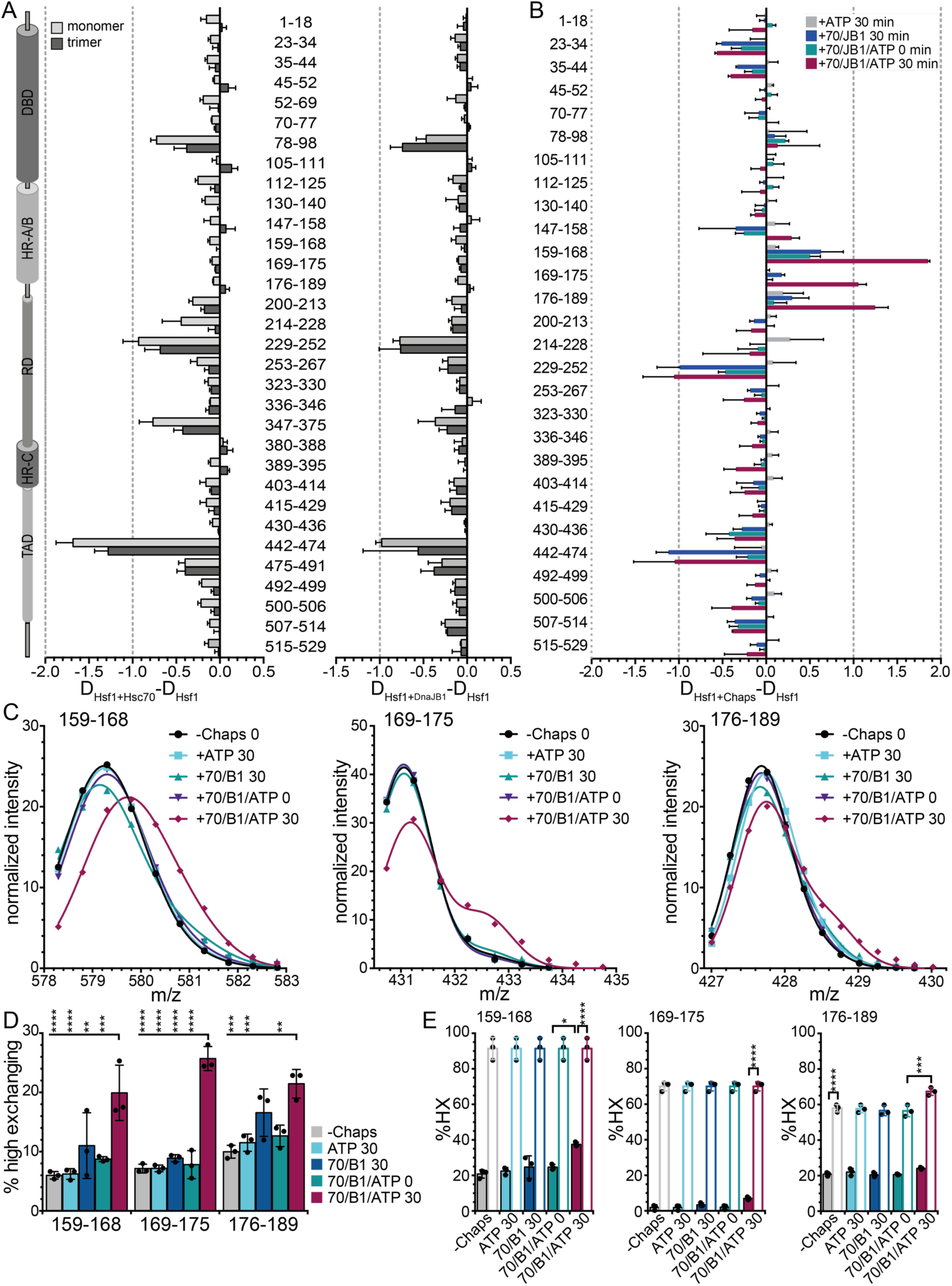
Hsc70 and DnaJB1 interaction with Hsf1 trimers lead to deprotection of segments within the trimerization domain. **A**, Difference plot of deuteron incorporation into Hsf1 monomers (light gray) or trimers (dark gray) in the presence of Hsc70 (left) or DnaJB1 (right) minus deuteron incorporation into Hsf1 in the absence of chaperones after 30 s incubation in D_2_O buffer. Shown are mean ± SD (n = 3) for peptic peptides as indicated between the panels. Left, Hsf1 domain representation. (see Fig. S2). **B**, Difference plot of deuteron incorporation into Hsf1 trimers incubated for 300 s in D_2_O buffer after pre-incubation in the presence of ATP, Hsc70 (70), and DnaJB1 (B1) as indicated minus deuteron incorporation into Hsf1 in the absences of additives. For relative deuteron incorporation see Fig. S4A. **C**, Fit of two Gaussian distributions to the peak intensities for peptic peptides 159-168 (left), 169-175 (middle), and 176-189 (right). Representative plots of 3 independent experiments. Original spectra and individual Gaussian distributions are shown in Fig. S4B and C. **D**, Fraction of high exchanging subpopulation for peptides 159-168, 169-175, and 176-189 under the different indicated pre-incubation conditions: -chaperones, +Hsc70 (70), +DnaJB1 (B1), and +ATP incubated for 0 or 30 min at 25°C. Statistical analysis: ANOVA with Sidak’s multiple comparison of different conditions for each of the peptides (n = 3); **, p < 0.01; ***, p < 0.001. **E**, Relative deuteron incorporation of the low (solid bars) and high (open bars) exchanging subpopulations. ANOVA, Sidak’s multiple comparison test; *, p < 0.05; ***, p < 0.001; ****, p < 0.0001.

Based on an Hsp70 binding site prediction algorithm, originally derived from peptide library scanning data for the *E. coli* Hsp70 homolog DnaK ^41^, two of the segments (200-213 and 442-474) protected from hydrogen exchange by Hsc70 covered sequences that fitted properties of strong Hsp70 binding sites (Fig. 4A, values below −5, dashed line). Close inspection of the sequence of these two segments revealed that both contained two potential Hsc70 binding sites.

**Figure 4:**
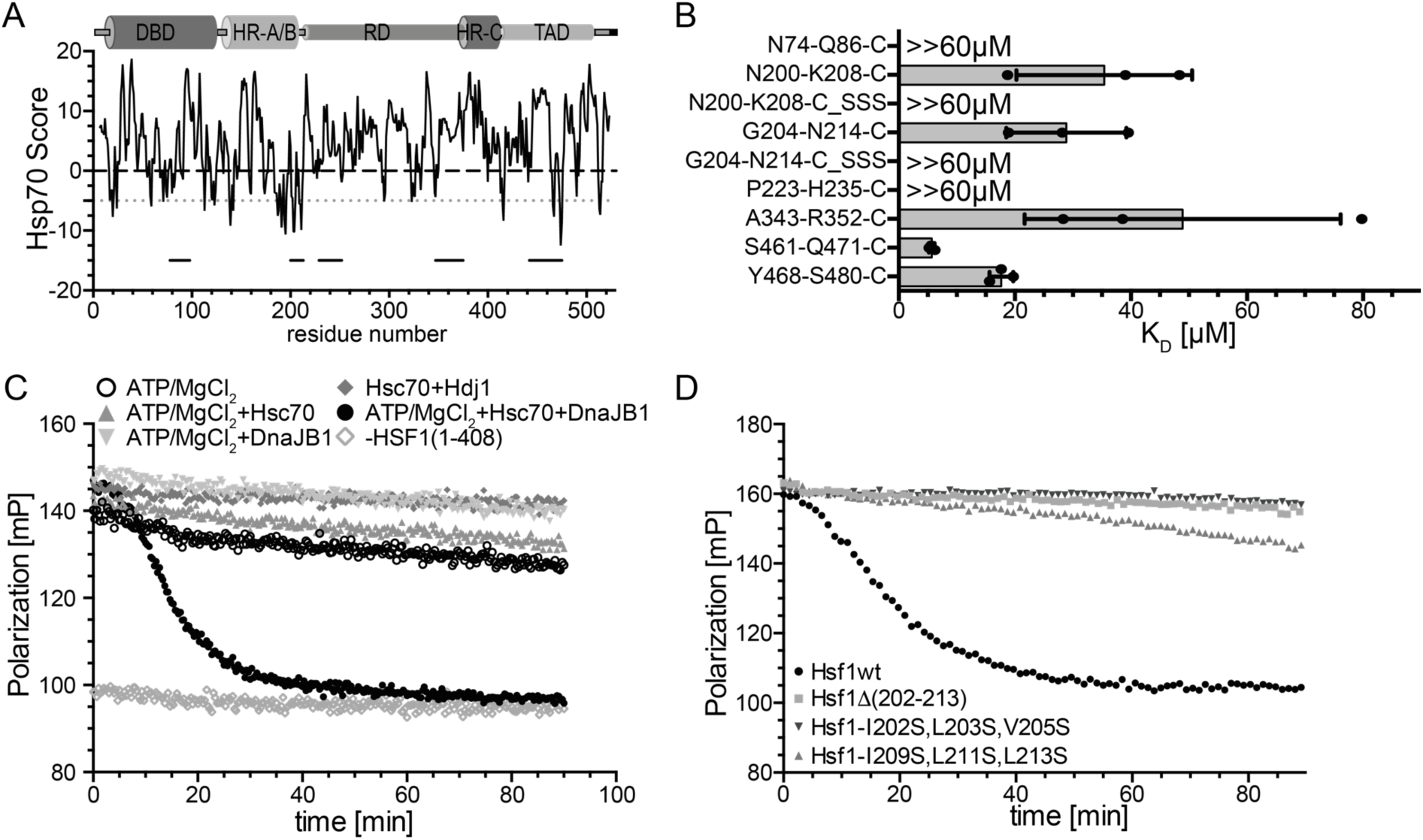
An Hsc70 binding site close to the trimerization domain of Hsf1 is necessary for Hsc70 mediated dissociation of Hsf1 from DNA. **A**, Hsp70 binding score as predicted by a published algorithm ^41^. The score value is attributed to the center of the sequence window of 13 residues. Values below −5 are considered to be good Hsp70 binding sites. Small horizontal lines represent the segments protected from hydrogen exchange. **B**, K_D_ values for the indicated peptides as determined by fluorescence polarization. **C**, Hsf1(1-408) with a deleted TAD is dissociated from HSE-DNA by Hsc70, DnaJB1 and ATP with similar rates as Hsf1wt. Fluorescence polarization assays containing Alexa 488-HSE-DNA bound Hsf1(1-408) trimers and the indicated components. **D**, The HR-B proximal Hsc70 binding site is essential for Hsc70/DnaJB1-mediated dissociation of HSE-DNA bound Hsf1.

To confirm Hsc70 binding to these sequences, we used peptides encompassing the respective sequences labeled with fluorescein and titrated Hsc70 concentration measuring fluorescence polarization. For all four potential binding sites we could determine a K_D_ between 5 and 30 µM (Fig. 4B). To the other protected segments Hsc70 did not have any measurable affinity. The highest affinity Hsc70 binding site was in the TAD, residues 461-471. All sites protected by Hsc70 were also protected by DnaJB1 consistent with the generally accepted mechanism of Hsp70 systems that J-domain proteins bind to the client protein first and target Hsp70 to its clients ^35^. When Hsc70, DnaJB1 and ATP were added to Hsf1 and pre-incubated for 30 min before dilution into D_2_O buffer and incubation for 300 s, we observed significant deprotection in three segments encompassing the trimerization domain (Fig. 3B, Fig. S3A), consistent with Hsc70-mediated monomerization. Close inspection of the respective spectra revealed a bimodal distribution of the isotopic peaks, indicative for the coexistence of two subpopulations with different exchange properties. Fitting the two Gaussian distributions to the isotope peak intensities allowed to extract the fraction of high and low exchanging subpopulations (Fig. 3C, Fig. S3B and C). In the absence of chaperones, in the presence of ATP, or in the presence of Hsc70, DnaJB1 but without ATP or with ATP but without pre-incubation only a small fraction of the peptides belonged to the high exchanging subpopulation (≤10% for 159-168 and 169-175; ≤17% for segment 176-189). Whereas upon pre-incubation for 30 min at 25°C in the presence of Hsc70, DnaJB1 and ATP the high exchanging subpopulation increased to 20 to 30%. In this case the high exchanging subpopulation exchanged 67 to 91% of the amide protons whereas the low exchanging subpopulation only exchanged 7 to 37% (Fig. 3E), indicating that in the presence of Hsc70, DnaJB1 and ATP the helices of the leucine-zipper are unfolded.

### Binding sites adjacent to HR-B are essential for Hsc70-mediated Hsf1 monomerization

To elucidate whether the Hsc70 binding sites in the TAD are responsible for Hsc70-mediated dissociation of Hsf1 from DNA and also to assess any involvement of the previously suggested Hsc70 binding site, we deleted the entire TAD, residues 409-529. Hsf1(1-408) was dissociated from DNA by Hsc70 with similar kinetics as wild type Hsf1 (Fig. 4C), excluding any involvement of sites in the TAD in this process. Consequently, we deleted the sites close to the HR-B region, residues 202-213. This deletion in Hsf1 completely prevented dissociation of Hsf1 from DNA in the presence of Hsc70, DnaJB1 and ATP, indicating that these residues are essential for Hsc70-mediated Hsf1 monomerization (Fig. 4D). Since a typical Hsc70 binding site consists of a core of up to five hydrophobic residues, preferably leucine, flanked by positively charged regions ^41^, we exchanged the hydrophobic residues against serine, a small hydrophilic residue disfavoring Hsc70 binding, generating the two Hsf1 variants, Hsf1-I202S,L203S,V205S and Hsf1-I209S,L211S,L213S. Hsc70-mediated dissociation of Hsf1 from DNA was either completely abrogated by these amino acid replacements or strongly inhibited (Fig. 4D). Peptides harboring the same amino acid replacements were also not bound by Hsc70 (Fig. 4B).

Taken together these data indicate that Hsc70-mediated dissociation of Hsf1 from DNA requires binding of Hsc70 to a site close to the trimerization domain. Hsc70 binding to the TAD may serve a different purpose.

### The number of available Hsc70 binding sites determines the rate of Hsf1 monomerization

We wondered whether binding of a single Hsc70 to one protomer of the Hsf1 trimer is sufficient for Hsc70-mediated monomerization. To address this question, we mixed monomeric Hsf1wt and Hsf1Δ(202-213) at different ratios and incubated the mixtures at 42°C for 10 min to form heterotrimers. Assuming that heterotrimers formed with the same probability as Hsf1wt and Hsf1Δ(202-213) homotrimers, different homo- and heterotrimeric species are formed according to a binomial distribution. At a 2:1 ratio of Hsf1wt:Hsf1Δ(202-213) 29.6% of the Hsf1 trimers are expected to be Hsf1wt homotrimers and consequently have three Hsc70 binding sites available close to the trimerized region, 44% are expected to have two Hsf1wt protomers per Hsf1 trimer, 22.2% are expected to have a single Hsf1wt per Hsf1 trimer, and 3.7% are expected Hsf1Δ(202-213) homotrimers. The fraction of the different homo- and heterotrimers change at the different mixing ratios as indicated in Fig. 5A. When we subjected these mixtures to Hsc70-mediated dissociation from DNA, we observed that the rate of dissociation decreased significantly with decreasing Hsf1wt:Hsf1Δ(202-213) ratios (Fig. 5A, Fig. S4B). Assuming that binding of a single Hsc70 is sufficient for Hsf1 monomerization and thus dissociation from DNA at wild type rates and that only the Hsf1Δ(202-213) homotrimer cannot be dissociated from DNA, as shown above, the overall rate of dissociation should not change for the different mixing ratios. Only the total amplitude of the dissociation reaction should change, because the Hsf1Δ(202-213) homotrimers would remain bound to a fraction of the DNA and thus cause a residual average polarization higher than the polarization of unbound DNA. We simulated this situation and found that the corresponding curves do not fit our experimental data (dashed lines in Fig. S4A). The alternative hypothesis that always three Hsc70 binding sites are necessary would lead to a similar situation, just with smaller amplitudes, also not explaining our experimental data. We concluded that the rate of Hsf1 monomerization depends on the number of Hsc70s bound simultaneously or sequential to different protomers of the Hsf1 trimer. An equation describing this situation fitted our experimental data reasonably well (Fig. S4C). Binding of a single Hsc70 per Hsf1 trimer is able to monomerize the Hsf1 trimer albeit at a very low rate, whereas action of Hsc70 on two protomers of the Hsf1 trimer more efficiently monomerizes Hsf1 trimers, and action of Hsc70 on all three protomers within the Hsf1 trimer is still more efficient in monomerization (Fig. S4D).

**Figure 5:**
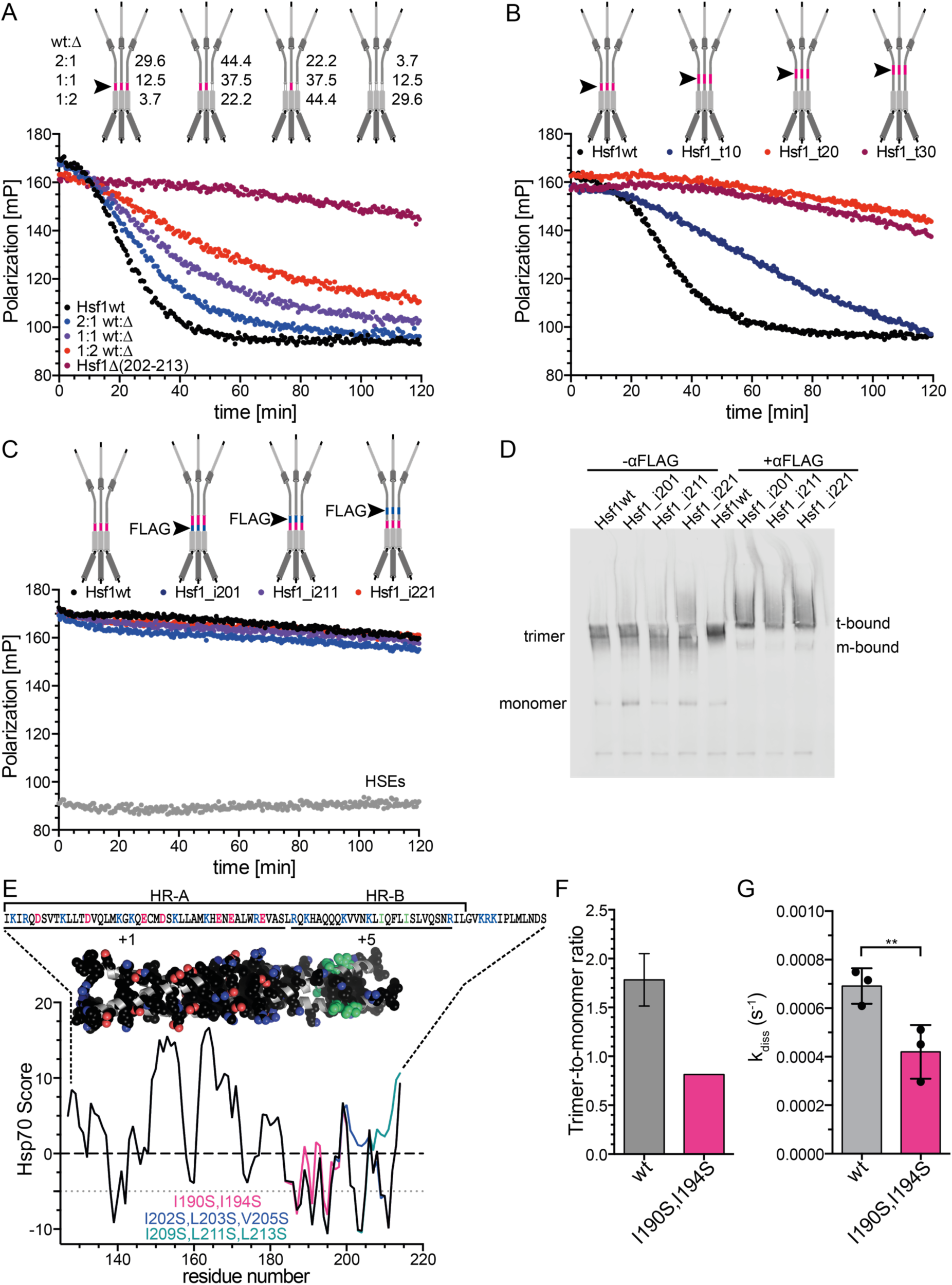
Hsc70/DnaJB1 monomerize Hsf1 by cycles of successive entropic pulling. **A**, The rate of Hsc70/DnaJB1-mediated dissociation of HSE-DNA bound Hsf1 depends on the number of HR-B proximal Hsc70 binding sites in the Hsf1 trimer. Hsf1wt (wt) and Hsf1Δ(202-213) (Δ) monomers were mixed in the indicated ratios and heat shocked at 42°C for 10 min to form mixtures of different homo- and heterotrimers as indicated with the cartoons (the red line and the arrow head indicate the Hsc70 binding site). Numbers indicate the relative fraction of the different species. For fit of the data see Fig. S4. **B**, Moving the Hsc70 binding site away from the trimerization domain reduces the rate of Hsc70/DnaJB1-mediated dissociation of HSE-DNA-bound Hsf1. Hsf1_t10/20/30, region 202-213 moved by 10, 20, or 30 residues towards the C-terminus. **C**, Anti-FLAG antibodies are not able to dissociate HSE-DNA-bound Hsf1. Red lines in the cartoon indicate the Hsc70 binding site; blue lines and arrow head indicate the inserted FLAG epitope DYKDDDDK. Hsf1_i201/211/221, FLAG-epitope inserted after residue 201, 211 or 221. Anti-FLAG antibodies were split in halfmers by incubation with 2 mM DTT (see Fig. S5). **D**, Anti-FLAG antibody halfmers bound to FLAG-epitope containing Hsf1 variants. Hsf1wt and FLAG-insertion variants were analyzed by blue-native gel electrophoresis in the absence or presence of DTT-treated anti-FLAG antibodies as indicated and Hsf1 detected by immunoblotting; t-bound, FLAG antibody-bound Hsf1 trimers; m-bound, FLAG antibody bound Hsf1 monomer. **E**, Predicted Hsc70 binding sites in the trimerization domain of Hsf1. Hsp70 score of the trimerization region residues 130-216 of wild type Hsf1 (black); Hsf1-I190S,I194S, magenta; Hsf1-I202S,L203S,V205S, blue; and Hsf1-I209S,L211S,L213S, green. Values below −5 are considered good Hsc70 binding sites. Above the graph is the trimeric homology model of the trimerization domain residues 130-203 with side chains of hydrophobic and charged residues in space filling representation in atom colors with carbon in black except for Ile190 and Ile194 where carbon is shown in green. Above the model is the corresponding sequence. Positively charged residues, blue; negatively charged residues, red. Lines below the sequence indicate two distinct regions with net charge +1 and +5. **F**, Trimer-to-monomer ratio of freshly purified Hsf1wt and Hsf1-I190S,I194S in the absence of heat shock, determined by gel filtration and BN-Gel (see Fig. S6). Hsf1wt, mean ± SD (n = 3). **G**, Rate of Hsc70/DnaJB1-mediated dissociation of HSE-DNA bound Hsf1wt and Hsf1-I190S,I194S; mean ± SD (n=3); **, P < 0.01; (paired t-test).

### Hsc70 monomerizes Hsf1 by entropic pulling

The concept of entropic pulling was proposed for Hsp70 action during import of polypeptides into mitochondria and for protein disaggregation ^42^. Briefly, Hsp70 binds to incoming polypeptides close to the membrane. Due to the excluded volume of the bulky Hsp70 the conformational freedom of the polypeptide is limited and thus the entropy low. Movement of the peptide into the mitochondrial matrix increases the distance of the polypeptide bound Hsp70 to the membrane, allowing for more conformational freedom of the polypeptide and thus increases the entropy. Since chemical reaction can be driven by increase in entropy as well as decrease of enthalpy, a force is generated that pulls the polypeptide into the mitochondrial matrix. De Los Rios et al. calculated a pulling force of around 10 to 20 pN that decrease with increasing length of the incoming polypeptide and will reach 0 pN once about 30 residues are imported. To drive further import a new Hsp70 needs to bind to the incoming polypeptide close to the membrane.

To test this hypothesis, we moved the Hsc70 binding site away from the HR-B region along the intrinsically disordered regulatory domain. Already when the Hsc70 binding site is 10 residues away from HR-B, Hsc70 dissociated Hsf1 from DNA with a significantly lower rate (Fig. 5B). At a distance of 20 residues Hsc70 was not anymore able to dissociate Hsf1 from the DNA, indicating that monomerization was not anymore possible. These results suggest that Hsc70 monomerizes Hsf1 trimers by entropic pulling.

To substantiate this hypothesis, we inserted a FLAG epitope between HR-B and the Hsc70 binding site or at 10 and 20 residues distance to HR-B. We treated anti-FLAG antibodies with DTT to split them in half (Fig. S5) and added them to DNA bound Hsf1. Surprisingly, we did not observe any dissociation of Hsf1 (Fig. 5C). This was not due to a failure of the FLAG-antibody halfmers to bind to the FLAG epitope containing Hsf1 trimers as demonstrated by BN-PAGE (Fig. 5D).

We hypothesized that pulling from a single site at the end of the trimerization domain may not be sufficient to unzip the entire domain, since the trimerization domain has a length of 75 residues and the entropic pulling force failed already when Hsc70 bound more than 20 residues away from the leucine-zipper. Close inspection of the HR-A/B region revealed that the sequence contains a large number of hydrophobic residues, as expected for a leucine-zipper, but unexpectedly the C-terminal part of the zipper (HR-B) contains 5 positively charged residues, which favor Hsc70 binding, and not a single negatively charged residue, which disfavor Hsc70 binding. Thus, this region of the trimerization domain contains several potential Hsc70 binding sites, as also evident from the Hsc70 binding site prediction (Fig. 5E). To compromise Hsc70 binding in this region is rather difficult, since replacing hydrophobic residues could disturb the leucine-zipper and prevent Hsf1 trimerization altogether. We used a model of the trimerization domain kindly provided by A. Bracher ^31^ which ended at residue 182 and used the iTASSER homology modeling software (https://zhanglab.ccmb.med.umich.edu/I-TASSER/; ^43, 44^) to extend the model to residue 203. In this model we discovered two isoleucine residues (I190 and I194) that did not point towards the zipper interface. Replacing these two isoleucines by serines only moderately reduced the propensity of this region to bind to Hsc70 (red line in Fig. 5E) as compared to the more drastic changes introduced by replacing I202, L203 and V205 (blue line) or I209, L211, and L213 (green line) by serines. Surprisingly, when we purified Hsf1-I190S,I194S we retrieved by gel filtration significantly less trimers and more monomers as compared to Hsf1wt (Fig. 5F), suggesting that the amino acid replacements have destabilized the trimeric state. Nevertheless, Hsc70-mediated monomerization and thus dissociation from DNA occurred at significantly lower rate for Hsf1-I190S,I194S than for Hsf1wt. These data suggest that binding of Hsc70 to this region contributes to Hsf1 monomerization. Altogether our results suggest that Hsc70 monomerizes Hsf1 by successive entropic pulling unzipping the leucine-zipper step by step.

### Amino acid replacements in Hsc70-binding sites potentiate heat shock reporter expression

To test the consequences of our findings in an *in-vivo*-model system, we stably transfected HSF1^-/-^ mouse embryonic fibroblasts (MEF) with a heat shock reporter expressing mTag-BFP under the control of the HSPA6 promoter. We then transiently transfected these cells with plasmids expressing wild type and mutant Hsf1 and rhGFP in an artificial operon using an IRES element (Fig. 6A) and analyzed the cells by flow cytometry, comparing the median BFP fluorescence level in GFP positive cells (Fig. 6B). Deletion or mutation of the HR-B proximal Hsc70 binding site increased BFP expression levels by up to fourfold and mutation in the Hsc70 binding site in the TAD increased BFP expression roughly twofold (Fig. 6C). In contrast, deletion or mutation of the previously proposed Hsc70 binding site in the TAD did not increase BFP expression. These data clearly demonstrate that binding of Hsc70 to the HR-B proximal Hsc70 binding site is important for attenuation of heat shock gene transcription and binding of Hsc70 to the newly identified Hsc70 binding site in the TAD contributes to attenuation presumably through interference of Hsf1 interaction with the core transcription machinery.

**Figure 6:**
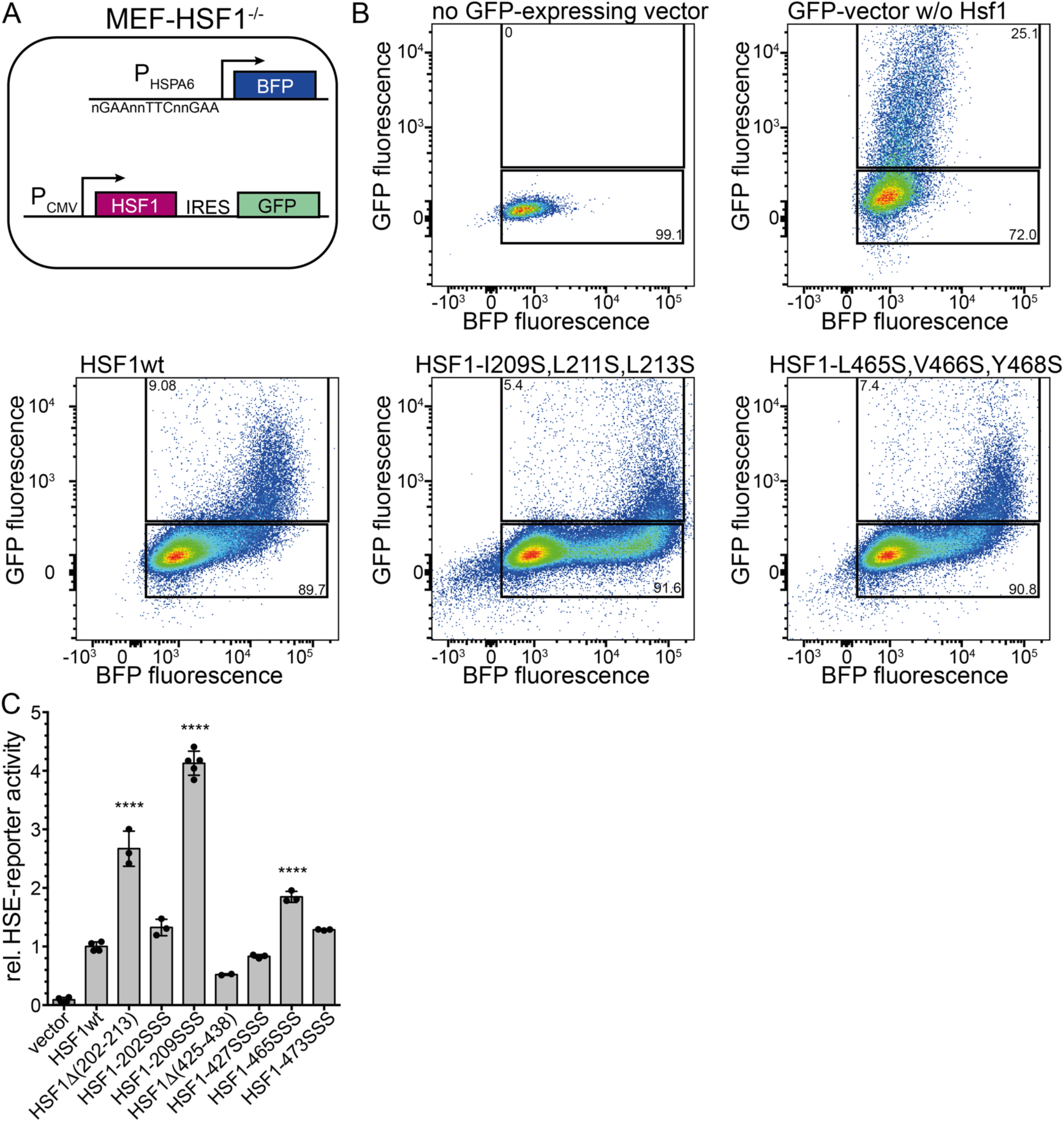
Compromising Hsc70 binding potentiates expression of a heat shock reporter. **A**, Schematics of the used system. HSF1^-/-^ MEFs were stably transfected with mTag-BFP expressed under the control of the HSPA6 promoter and subsequently transfected with plasmids that expressed HSF1wt and mutant variants in an artificial operon with rhGFP, which is translated from an IRES element. **B**, Exemplary flow cytometry data plotting GFP fluorescence versus BFP fluorescence, indicating the gates used for analysis. Numbers indicate the fraction of cells in each gate. **C**, Median of BFP fluorescence of GFP positive cells minus median of BFP fluorescence of GFP negative cells. Median ±SD of at least 2 independent experiments.

## Discussion

In this study we gained several important insights into the regulation of the HSR. We demonstrate that Hsc70 together with its J-domain co-chaperone DnaJB1 removes Hsf1 from heat shock promoter DNA by monomerizing the Hsf1 trimers. Thus, the HSR can be shut off without HSF1 acetylation or degradation and HSF1 can be recycled. Hsf1 monomerization starts from a Hsc70 binding site C-terminal of the trimerization domain proximal to HR-B and proceeds towards the N-terminus of the trimerization domain through stepwise unzipping the leucine-zipper by entropic pulling (Fig. 7A). We show that starting this repeated binding of Hsc70 at several protomers of the Hsf1 trimer allows for more rapid disassembly. This mechanism makes the biggest contribution to attenuation of heat shock gene transcription in a cell culture model. We also found that binding of Hsc70 to a newly identified site in the TAD contributes to attenuation in the cell culture model, most likely by interfering with the Hsf1-mediated release of stalled RNA polymerase. It is interesting that this binding site has the highest affinity for Hsc70 (K_D_ ca. 5 µM), indicating that increasing concentrations of free Hsc70 during heat shock transcription will on average first bind to this site before binding to the HR-B proximal site.

**Figure 7:**
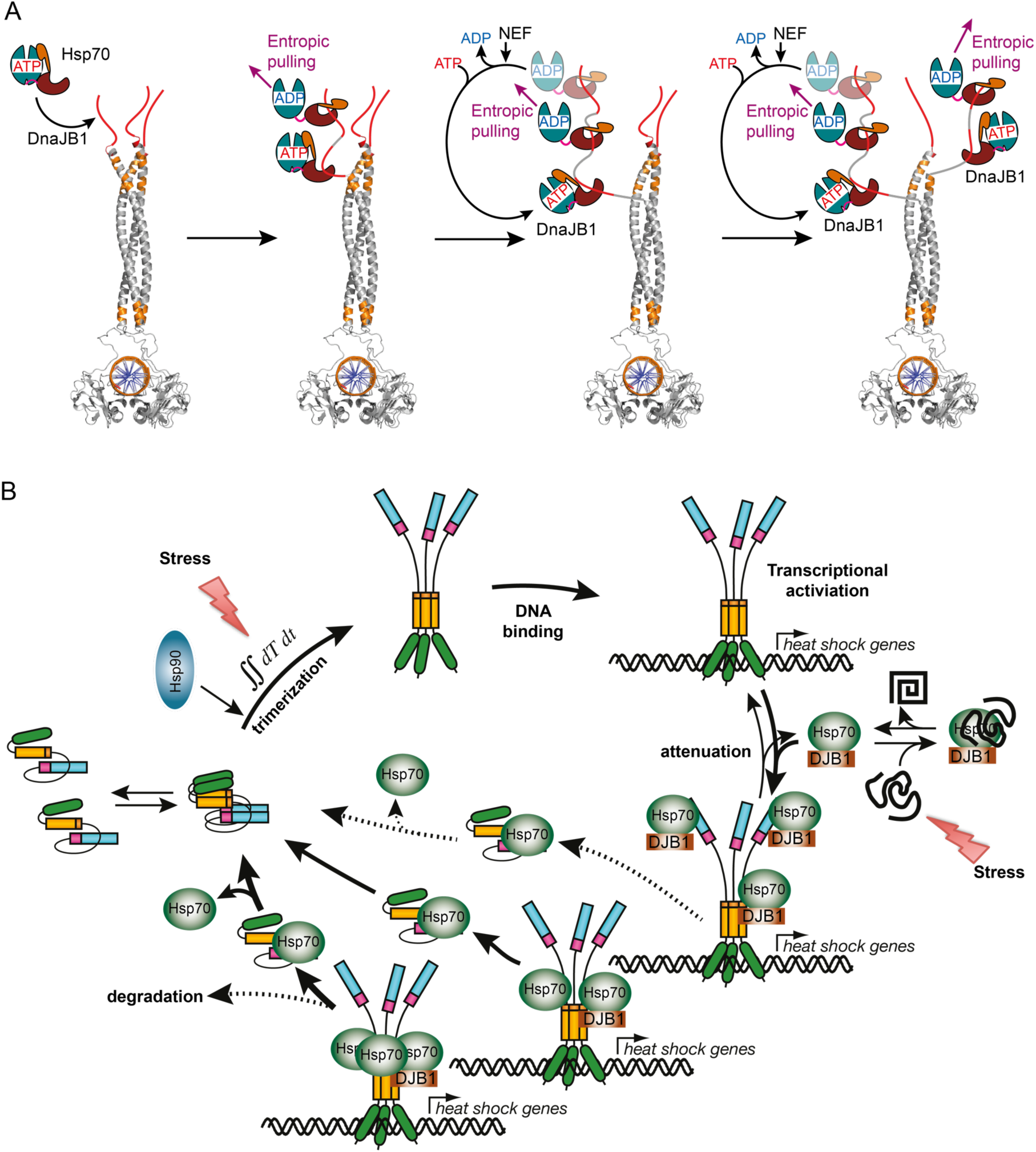
Model of the Hsc70 mediated regulation of the heat shock response. **A**, Illustration of the stepwise disassembly of Hsf1 trimers by the entropic pulling action of Hsc70 and DnaJB1. Homology model of trimeric DNA bound human Hsf1. Red, accessible Hsc70 binding sites; orange, Hsc70 binding sites not accessible in the Hsf1 trimer, only accessible after unfolding. Unzipping of the Hsf1 trimer may occur simultaneously on all three protomers. **B**, Hsf1 activation-attenuation cycle. In unstressed cells Hsf1 is in monomer-dimer equilibrium, occasionally trimerizing depending on the local concentration. Trimeric Hsf1 either binds to HSE-promoter DNA driving heat shock gene transcription by releasing paused RNA polymerase or is disassembled immediately by Hsc70/DnaJB1 action (not shown for clarity). Hsc70 binds dynamically to the TAD of DNA-bound Hsf1 trimers attenuating transcriptional activity and bind to its HR-B proximal binding site, disassembling Hsf1 trimers and thus removing it from heat shock promoters. Release of Hsf1 from Hsc70 restarts this cycle. Upon heat shock Hsf1 trimerizes at elevated rates due to its thermosensory function, integrating over temperature and time at elevated temperatures. Simultaneously, Hsc70/DnaJB1-mediated attenuation and disassembly is slowed down, due to binding of Hsc70 to misfolded and aggregated proteins. Both parts of the cycle shift the Hsf1 pool rapidly to the DNA bound active state, accelerating heat shock gene transcription. Elevating Hsc70 concentrations successively attenuate transcriptional activity of Hsf1 and disassemble Hsf1 trimers.

One might wonder why the affinity of Hsc70 for the critical HR-B proximal site is so low (K_D_ ca. 30 µM). Firstly, it is important to understand that the equilibrium dissociation constant determined for the ADP bound state only insufficiently describes the real affinity of Hsc70 for a binding site. In the presence of ATP and a J-domain protein, Hsc70·ATP associates with high rates with the protein client, then the J-domain protein in synergism with the substrate polypeptide stimulates ATP hydrolysis in Hsc70, leading to transition to the so-called high affinity state with low substrate dissociation rates. This targeting mechanism leads to a non-equilibrium situation that decreases the apparent K_D_ by several orders of magnitude and was coined ultra-affinity ^45^. The actual affinity of Hsc70 to this site depends on the local concentration of DnaJB1, which seems to have also some affinity for these sites to increase local concentration and the concentration of the nucleotide exchange factor (Fig. 1I) that allows for ADP release, ATP rebinding, and substrate release. Secondly, the cellular concentration of Hsc70 averaged over 11 different cancer cell lines is around 12-18 µM under non-stress conditions ^46^ assuming a total cellular protein concentration of 100 – 150 mg/ml ^47, 48^ and can reach 14 to 21 µM upon a mild heat shock of 41°C for 4 h ^48^.

In contrast, human Hsp70 (HSPA1A/B) is barely detectable in most non-cancer cells and reaches only about 0.04 to 0.06 µM upon heat shock ^48^, but can reach around 0.7 – 10 µM in cancer cells ^46^. It is very likely that the HR-B proximal site evolved for such an affinity to Hsc70 to allow for a high enough concentration of Hsc70 in the cell. If the affinity of this site for Hsc70 were higher, Hsc70 might disassemble Hsf1 already at lower concentrations when the cell still needs more Hsc70 and other stress proteins. This also explains why the reaction is so exquisitely sensitive to the concentration of Hsc70 and DnaJB1 and why high concentrations of the nucleotide exchange factor Apg2 inhibit the reaction. Apg2 accelerates nucleotide exchange and thus release of Hsc70 from Hsf1. If the first Hsc70 is released from Hsf1 before another Hsc70 can bind to the next Hsc70 binding site that becomes transiently accessible through entropic pulling at the first site, unzipping cannot be efficient. Our data also demonstrate that working on several protomers of the Hsf1 trimer more efficiently disassembles Hsf1 trimers than pulling on a single protomer, providing an additional explanation for the necessity of higher Hsc70 concentrations. A reasonable explanation for this observation is that the entropic pulling force is larger, if several Hsc70 molecules are bound in close proximity to each other to individual protomers of the Hsf1 trimer, increasing local crowding and thereby decreasing the conformational freedom of the HR-B proximal region of the intrinsically disordered regulatory domain. It is interesting that inserting a FLAG epitope before or after the HR-B proximal Hsc70 binding site and using anti-FLAG antibody halfmers did not lead to disassembly of Hsf1 trimers. This observation contrasts disassembly of clathrin coats that was efficiently performed by replacing the Hsc70 binding site and using FLAG Fab fragments ^49^, indicating that a single pull at all three Hsf1 protomers in the Hsf1 trimer is not sufficient for disassembly. It would not be possible to add more FLAG epitopes along the trimerization domain because they would disrupt the triple leucine-zipper. Conceptionally, disassembly of Hsf1 trimers is more similar to protein translocation through a membrane and to disaggregation. In both cases successive binding events along a polypeptide chain pulls the protein through the membrane pore and out of the aggregate, respectively.

Based on the here presented data and published literature, we propose the following model for the regulation of the HSR (Fig. 7B). In unstressed cells Hsf1 is in a monomer-dimer equilibrium occasionally trimerizing in a concentration and Hsp90-dependent manner and bind to HSEs in the genome. Hsc70 may bind to the TAD to attenuate heat shock gene transcription and to the HR-B proximal Hsc70 binding site to disassemble either free Hsf1 trimers before DNA binding (not shown in Fig. 7B) or DNA bound Hsf1 trimers into monomers. This situation leads to a low constitutive transcription of heat shock genes. Upon heat shock Hsf1 trimerization is accelerated by Hsf1 thermosensing, integrating over temperature and time spent at elevated temperatures ^10^. At the same time Hsf1 monomerization is slowed down, due to titration of Hsc70 to misfolded and aggregated proteins. Thus, heat shock gene transcription increases rapidly and Hsc70 and Hsp70 concentrations increase. As soon as the concentration of free Hsc70/Hsp70 are sufficient Hsc70 or Hsp70 bind to the TAD of Hsf1, the site with the highest affinity, reducing transcriptional activation. This interaction is transient following the known Hsp70 ATPase cycle and depend on the nucleotide exchange factor concentration. As Hsc70 and Hsp70 concentrations continue to increase, Hsc70/Hsp70 concentrations will be sufficient to bind to the HR-B site and to disassemble the Hsf1 trimer and thus removing it from DNA, reducing heat shock gene transcription to a level that keeps Hsc70 and Hsp70 at the necessary concentrations. This Hsf1 activation/attenuation cycle is highly sensitive to the concentration of free Hsc70. Our model explains a host of literature accumulated over the last 40 years. This cycle may be modulated at many sites for example by abundance regulation of Hsf1, by nuclear-cytoplasmic transport of Hsf1 and Hsc70 or nucleotide exchange factors like Apg2, and by posttranslational modifications that may influence binding of Hsc70 to Hsf1. In particular, phosphorylation which was shown to accompany heat shock induction and to prolong heat shock gene transcription is expected to interfere with Hsc70 binding due to the fact that negative charges disfavor Hsp70 binding ^41^.

## Acknowledgement

We thank Dr. G. Stöcklin for providing the HSF1^-/-^ MEFs, Dr. M. Langlotz for help in the FACS facility, Dr. T. Ruppert and N. Lübbehusen for help in the core facility for mass spectrometry and proteomics, and S. Hennes und A. Müller for excellent technical assistance. This work was funded by the Deutsche Forschungsgemeinschaft (SFB 1036 TP9).

## Authors’ contributions

Conceptualization, M.P.M.; Methodology, L.L-B., S.W.K., M.P.M.; Investigation, L.L-B., S.W.K., and M.P.M.; Writing – Original Draft, S.W.K. and M.P.M.; Writing – Review & Editing, M.P.M., S.W.K. and L.L-B.; Funding Acquisition, M.P.M.; Supervision, M.P.M.

## Methods

### Protein Expression and Purification

Human Hsf1, Hsf1(1-408), Hsf1Δ(202-213), Hsf1-I202S,L203S,V205S, Hsf1-I209S,L211S,L213S, Hsf1ΔT(202-213), Hsf1ΔT(202-213), Hsf1ΔT(202-213), Hsf1_i201(DYKDDDDK), Hsf1_i211(DYKDDDDK) or Hsf1_i221(DYKDDDDK) were purified as N-terminal His_6_-SUMO fusions from *E. coli* BL21 (DE3) Rosetta cells after overproduction at 20°C for 2 h. Bacterial pellets were resuspended in lysis buffer (25 mM HEPES/KOH pH 7.5, 100 mM KCl, 5 mM MgCl_2_ and 10% glycerol), drop-wise frozen in liquid nitrogen, disrupted using a Mixer Mill MM400 (Retsch) and resuspended in 200 ml Hsf1 lysis buffer supplemented with 3 mM β-mercaptoethanol, DNase and protease inhibitors (10 μg/ml aprotinin, 10 μg/ml leupeptin, 8 μg/ml pepstatin, 1 mM PMSF). The resulting lysate was centrifuged (33000 x g, 4°C for 45 min) and the soluble fraction incubated for 25 min at 4°C with 0.4 g Protino Ni-IDA resin (Macherey-Nagel) in a rotation shaker. The resin was transferred to a gravity-flow column and washed with 100 ml of Hsf1 lysis buffer. Bound protein was eluted with Hsf1 lysis buffer containing 250 mM imidazole, 3 mM β-mercaptoethanol and protease inhibitors. The His_6_-SUMO tag was cleaved off by incubation with Ulp1 SUMO-protease for 1 h at 4°C. The cleaved Hsf1 was further separated by size-exclusion on Superdex 200 HiLoad 16/60 column (GE Healthcare), equilibrated with Hsf1 SEC buffer (25 mM HEPES/NaOH pH 7.5, 150 mM NaCl, 10% glycerol, 2 mM DTT). The fractions containing either monomeric or trimeric Hsf1 were concentrated to 10 μM, flash-frozen in liquid nitrogen and stored at −80°C.

Human **DnaJB1/Hdj1** and DNAJB1-H32Q,D34N were purified as a His_6_-SUMO fusion from *E. coli* BL21 (DE3) Rosetta cells after overproduction for 3 h at 30°C. Cells were resuspended in DnaJB1 lysis buffer (50 mM HEPES/KOH pH 7.5, 750 mM KCl, 5 mM MgCl_2_, 10% glycerol) supplemented with 3 mM β-mercaptoethanol, DNase and protease inhibitors (10 μg/ml aprotinin, 10 μg/ml leupeptin, 8 μg/ml pepstatin, 1 mM PMSF). Bacteria were lysed using chilled microfluidizer at a pressure of 1000 bar. The resulting lysate was centrifuged (33000 x g, 4°C for 45 min) and the supernatant incubated for 25 min at 4°C with 1.5 g Protino Ni-IDA resin (Macherey-Nagel) in a rotation shaker. The resin was transferred to a gravity-flow column, washed with 200 ml of DnaJB1 lysis buffer and then with 100 ml of DNaJB1 SEC buffer (50 mM HEPES/KOH pH 7.5, 500 mM KCl, 5 mM MgCl_2_, 10% glycerol). Protein was eluted with DnaJB1 SEC buffer containing 300 mM imidazole, 3 mM β-mercaptoethanol and protease inhibitors. Fractions containing DnaJB1 were desalted (HiPrep 26/10 Desalting column, GE Healthcare) to SEC buffer and digested with Ulp1 protease overnight. After proteolytic cleavage the protein was incubated with 1g Protino Ni-IDA resin (Macherey-Nagel) for 25 min at 4°C to remove His_6_-SUMO. The flow through was collected and further purified by size-exclusion on Superdex 200 HiLoad 16/60 column (GE Healthcare) equilibrated with DnaJB1 SEC buffer.

Human **Hsc70 (HSPA8), Hsc70-K71M, Hsc70-V438F and Hsp70 (HSPA1A)** were purified as a His_6_-SUMO fusion from overproducing *E. coli* BL21 (DE3) Rosetta. Cells were resuspended in Hsc70 lysis buffer (50 mM Tris pH 7.5, 300 mM NaCl, 5 mM MgCl_2_, Saccharose 10%) supplemented with 3 mM β-mercaptoethanol, DNase and protease inhibitors (10 μg/ml aprotinin, 10 μg/ml leupeptin, 8 μg/ml pepstatin, 1 mM PMSF). Bacteria were lysed using a chilled microfluidizer at a pressure of 1000 bar. The resulting lysate was centrifuged (33000 x g, 4°C for 45 min) and the supernatant was incubated for 25 min at 4°C with 2 g Protino Ni-IDA resin (Macherey-Nagel) in a rotation shaker. The resin was transferred to a gravity-flow column, washed with 200 ml of Hsc70 lysis buffer followed by high salt (Hsc70 lysis buffer but 1 M NaCl) and ATP (Hsc70 lysis buffer with 5 mM ATP) washes. Hsc70/Hsp70 were eluted with Hsc70 lysis buffer containing 300 mM imidazole, 3 mM β-mercaptoethanol and protease inhibitors. Fractions containing Hsc70 were desalted (HiPrep 26/10 Desalting column, GE Healthcare) to HKM150 buffer (25 mM HEPES/KOH pH 7.6, 150mM KCl, 5mM MgCl_2_) and digested with Ulp1 protease overnight. After proteolytic cleavage the protein was incubated with 1.5 g Protino Ni-IDA resin (Macherey-Nagel) for 25 min at 4°C to remove His_6_-SUMO. Flow through was collected, desalted to HKM150 buffer, concentrated to 50 μM, aliquoted, flash-frozen in liquid nitrogen and stored at −80°C.

Human **Hsp90**α, Hsp90α-E47A were purified as a His_6_-SUMO fusion from *E. coli* BL21 (DE3) Rosetta cells after overproduction overnight at 25°C. Cells were harvested by centrifugation (5000 x g, 4°C for 10 min), resuspended in Hsp90α lysis buffer (20 mM HEPES/KOH pH 7.5, 100 mM KCl, 5 mM MgCl_2_, 10% glycerol) supplemented with 3 mM β-mercaptoethanol, DNase and protease inhibitors (10 μg/ml aprotinin, 10 μg/ml leupeptin, 8 μg/ml pepstatin, 1 mM PMSF), and lysed using a chilled microfluidizer at a pressure of 1000 bar. The resulting lysate was centrifuged (33,000 x g, 4°C for 45 min) and the supernatant was incubated for 25 min at 4°C with 1.5 g Protino Ni-IDA resin (Macherey-Nagel) in a rotation shaker. The resin was transferred to a gravity-flow column and washed with 200 ml of Hsp90α lysis buffer. Protein was eluted with Hsp90α lysis buffer containing 400 mM imidazole, 3 mM β-mercaptoethanol and protease inhibitors. Fractions containing Hsp90α were subjected to Ulp1 cleavage and desalting during overnight dialysis to Hsp90α dialysis buffer (20 mM HEPES/KOH, pH 7.5, 20 mM KCl, 5 mM MgCl_2_, 10% glycerol, 3 mM β-mercaptoethanol). After proteolytic cleavage protein was incubated with 1.2 g Protino Ni-IDA resin (Macherey-Nagel) for 25 min at 4°C to remove His_6_-SUMO. The flow-through was subjected to anion exchange chromatography (ResourceQ column, GE Healthcare, 20 mM HEPES/KOH pH 7.5, 5 mM MgCl_2_, 10% glycerol, 20-1000 mM KCl gradient in 16 CV) followed by size-exclusion on Superdex 200® HiLoad 16/60 column (GE Healthcare) equilibrated with Hsp90α SEC buffer (40 mM HEPES/KOH, pH 7.5, 50 mM KCl, 5 mM MgCl_2_, 10% glycerol, 3 mM β-mercaptoethanol). The fractions containing Hsp90α were pooled, concentrated to 50 μM, aliquoted, flash-frozen in liquid nitrogen and stored at −80°C.

Human **Apg2** was purified as a His_6_-SUMO fusion from *E. coli* BL21 (DE3) Rosetta cells after overproduction overnight at 30°C. Cells were subsequently harvested by centrifugation (5000 x g, 4°C for 10 min), resuspended in Apg2 lysis buffer (40 mM Tris pH 7.9, 100 mM KCl, 5 mM ATP) supplemented with 3 mM β-mercaptoethanol, DNase and protease inhibitors (10 μg/ml aprotinin, 10 μg/ml leupeptin, 8 μg/ml pepstatin, 1 mM PMSF), and lysed using a chilled microfluidizer at a pressure of 1000 bar. The lysate was centrifuged (33000 x g, 4°C for 45 min) and the supernatant was incubated for 25 min at 4°C with 2 g Protino Ni-IDA resin (Macherey-Nagel) in a rotation shaker. The resin was transferred to a gravity-flow column and washed with 200 ml of Apg2 lysis buffer. Protein was eluted with Apg2 lysis buffer containing 300 mM imidazole, 3 mM β-mercaptoethanol and protease inhibitors. Fractions containing Apg2 were subjected to desalting to Apg2 desalting buffer (40 mM Tris pH 7.9, 100 mM KCl,). Desalted protein was supplemented with 10% glycerol, 5 mM ATP, protein inhibitors, cleaved overnight with Ulp1 proteaseand subsequently incubated with 1.5 g Protino Ni-IDA resin (Macherey-Nagel) for 25 min at 4°C to remove His_6_-SUMO. The flow-through was subjected to size-exclusion on Superdex 200 HiLoad 16/60 column (GE Healthcare) equilibrated with Apg2 SEC buffer (40 mM HEPES/KOH pH 7.6, 10 mM KCl, 5 mM MgCl_2_, 10% glycerol) followed by IEC (ResourceQ column, 40 mM HEPES/KOH pH 7.6, 5 mM MgCl_2_, 10% glycerol, 10-1000 mM KCl gradient in 10 CV). The fractions containing Apg2 were pooled, supplemented with 5 mM ATP, aliquoted, flash-frozen in liquid nitrogen and stored at −80°C.

### Blue native PAGE (BN-PAGE)

Protein samples pre-mixed with 4x BN-PAGE sample buffer (250 mM Tris, pH 6.8, 40% glycerol, 0.1% Coomassie Brilliant Blue G-250) were separated on 8% PAGE gels in Laemmli system (cold BN-PAGE running buffer: 25 mM Tris, 0.2 M glycine).

Separation was carried out at constant current (20 mA per gel, 1 hr, on ice). Gels were subsequently stained using staining solution (Quick Coomassie Stain, Serva) or used further for Western blot analysis; primary antibody (HSF1 (H-311) rabbit polyclonal IgG, Santa Cruz Biotechnology, 1:1000 dilution), secondary fluorescently labelled antibody (Goat anti rabbit IgG IRDye 680RD, Odyssay, 1:20000 dilution). Fluorescence was detected on LICOR Odyssey Infrared Imaging System (700 nm channel) and quantified using Image Studio Lite Ver 5.2.

### Fluorescence Spectroscopy

#### Binding of trimeric Hsf1 to Heat Shock Elements (HSEs)

Fluorescently labelled double stranded 3xHSEs were prepared by annealing of fluorescently labelled Alexa488-3xHSE sense oligonucleotides with 3xHSE antisense nucleotides (2 min at 70°C, 0.6°C/min stepwise decrease from 70°C to 30°C). 5 nM 3xHSEs were titrated with trimeric Hsf1 at different concentration range (from 0.2 nM to 100 nM). Protein was mixed with 3xHSEs at 1:1 ratio on low volume 384-well plate (CORNING, REF3820). The plate was subsequently spun down at 1000 x g for 1 min at RT. Fluorescence anisotropy of prepared samples was measured at 25°C using plate reader (CLARIOstar, BMG Labtech, Excitation, F:482-16, Emission, F:530-40). The data points were fitted to the quadratic solution of the law of mass action using GraphPad Prism software.

#### Hsf1 dissociation from Heat Shock Elements (HSEs)

Fluorescently labelled double stranded 3xHSEs were prepared by annealing of fluorescently labelled Alexa488-3xHSE sense oligonucleotides with 3xHSE antisense nucleotides (2 min at 70°C, 0.6°C/min stepwise decrease to 30°C). 5 μM Hsc70, 2.5 μM Hdj1/DNAJB1, 10 μM Hsp90α, 2 mM ATP, 4 mM MgCl_2_ in Hsf1 SEC buffer (25 mM HEPES/NaOH pH 7.5, 150 mM NaCl, 10% glycerol, 2 mM DTT) preincubated at 30°C for 30 min were mixed with 100 nM trimeric Hsf1 and 25 nM HSEs on low volume 384-well plate (CORNING, REF3820) in a final 20 μl reaction volume. The plate was subsequently spun down at 1000 x g for 1 min at RT. Fluorescence anisotropy of prepared samples was measured at 25°C using a microplate reader (CLARIOstar, BMG Labtech, Excitation, F:482-16, Emission, F:530-40). The trimeric Hsf1 fraction of the gelfiltration was used for the experiments, if not stated otherwise in the figure legend. A single exponential equation with delay was fitted to the data using Prism (GraphPad software) (see Fig. S1C).

#### Hsf1 FLAG-variants dissociation from HSEs

Anti-FLAG antibody (SIGMA) was desalted to Hsf1 SEC buffer (25 mM HEPES/NaOH pH 7.5, 150 mM NaCl, 10% glycerol, 2 mM DTT) and incubated for 1 h at 25°C to reduce it. Reduced antibody was used in the dissociation experiment (described above) at 5 μM final concentration. *Hsc70 binding to fluorescently labelled peptides.* Peptides were incubated with equimolar amounts of TCEP for 30 min at 25°C to reduce the C-terminal cysteine residue. 100 μM of peptide were mixed with 22-fold molar excess of fluorescein-5-maleimide (ThermoFischer SCIENTIFIC) in 25 mM HEPES/KOH pH 7.2 and incubated for 2 h at 25°C. Labelled peptides were separated from excess free dye on a Sephadex G-10 gravity column (GE Healthcare, 17-0010-01) equilibrated with HKM50 buffer (25 mM HEPES/KOH pH 7.6, 50 mM KCl, 5 mM MgCl_2_). Labelled peptides at 1 μM final concentration were titrated with Hsc70 (0-60 µM final concentration range) and incubated on low volume 384-well plate (CORNING, REF3820) in 20 μl final volume for two hours at 25 °C. Fluorescence anisotropy was measured using a CLARIOstar plate reader (BMG Labtech, Excitation, F:482-16, Emission, F:530-40). To avoid the formation of oligomeric species, Hsc70 was concentrated in the presence of 1 mM ATP. HKM150 (25 mM HEPES/KOH pH 7.6, 150 mM KCl, 5 mM MgCl_2_) was used as assay buffer. The quadratic solution of the law of mass action was fitted to the data using GraphPad Prism.

### Hsf1 trimers formation

Hsf1 can be purified as a trimer and monomer. Monomeric Hsf1 can be converted into trimeric Hsf1 by incubation at 42°C for 10 min. In case of Hsf1 heterotrimer formation, monomers of Hsf1wt and Hsf1Δ(202-213) were pre-mixed at different ratios and subjected to a 10-min-incubation at 42°C.

### Hydrogen-Exchange Mass Spectrometry - Sample preparation

Effects *of Hsc70 or Hdj1/DNAJB1 on Hsf1.* 6 μM Hsf1 (monomer or trimer) were pre-incubated for 30 min at 25°C in the presence of 12 μM Hsc70 or 9 μM Hdj1 in Hsf1 SEC buffer (25 mM HEPES/NaOH pH 7.5, 150 mM NaCl, 10% glycerol, 2 mM DTT) and, subsequently, diluted 10-fold in H_2_O/D_2_O buffer to a final volume of 100 μl and incubated for 30 s at 25°C. Deuteration reaction was quenched by adding 100 μl of ice-cold quench buffer (400 mM Na-phosphate, pH 2.2) and was immediately injected into the ice-cold HPLC system.

*Effects of the Hsp70 system on Hsf1.* 25 μM Hsf1 trimers were pre-incubated in the presence of 40 μM Hsc70, 12 μM Hdj1, 2 mM ATP, 4 mM MgCl_2_ in Hsf1 SEC buffer (25 mM HEPES/NaOH pH 7.5, 150 mM NaCl, 10% glycerol, 2 mM DTT) for 30 min at 25°C, subsequently, diluted 10-fold in D_2_O buffer to a final volume of 100 μl, and incubated for 300 s at 25°C. The deuteration reaction was quenched by adding 100 μl of the ice-cold quench buffer (2% formic acid) and was immediately injected into the ice-cold HPLC system.

### Performing Hydrogen-Exchange Mass Spectrometry measurements

In the ice cooled HPLC set-up the protein was digested online using a column with immobilized pepsin and the peptides were desalted on a C8 trap column (POROS 10 R2, Applied Biosystems, #1-1118-02) for 2 min and eluted over an analytical C8 column (Waters GmbH, 186002876) using a 10 min gradient from 5 to 55% acetonitrile. All experiments were performed using a Maxis mass spectrometer (Bruker, Bremen, Germany) and analyzed with the Data Analysis software. The calculated centroid values were corrected for the back-exchange using a 100% deuterated sample.

### Cell culture experiments

HSF1^-/-^ MEF (mouse embryonic fibroblasts) were a kind gift of Prof. I.J. Benjamin (University of Utah, School of Medicine, Salt Lake City Utah, USA) ^50^. The cells were cultured in high-glucose Dulbecco’s modified Eagle media (GIBCO, Thermo Fisher Scientific) with 10% fetal bovine serum (GIBCO, Thermo Fisher Scientific) 1% Penicilin/Streptomycin (Pen/Strep, Merck, Germany), and 1% minimum essential Eagle’s medium (Merck, Germany) at 37°C and 5% CO_2_. Stable and transient transfection was performed using TRansIT^®^-X2 Dynamic Delivery system (Mirus, Madison, USA) according to the manufactures instructions. HSF1^-/-^ MEF PHSPA6-mTagBFP cells were constructed by transfection of HSF1^-/-^ MEF cells with pcDNA5-ble-PHSPA6-mTagBFP, selection in DEMEM supplemented with Zeocin (InvivoGen 900 µg/ml) for 7-10 days and subsequently sorting by fluorescence activated cell sorting (FACS) on a BD FACS Aria III (BD Biosciences, excitation 407 nm, emission filter 450/40 nm) for BFP positive single cells. Individual clones were tested for successful integration of the Hsf1 activity-reporter by transient transfection with pIRES-HSF1_rhGFP.

To test HSF1 activity of Hsf1 wild type and mutant proteins, HSF1^-/-^ MEF PHSPA6-mTagBFP cells were transiently transfected with plasmids pIRES-HSF1_rhGFP, pIRES-HSF1Δ(202-213)_rhGFP, pIRES-HSF1-I202S,L203S,V205S_rhGFP, pIRES-HSF1-I209S,L211S,L213S_rhGFP, pIRES-HSF1L465S,V466S,Y468S_rhGFP, pIRES-HSF1-L473S,F474S,L475S,L476S_rhGFP, pIRES-HSF1Δ(425-438)_rhGFP, pIRES-HSF1-M427S,L249S,L432S,L436S_rhGFP, grown for 48 h at 37°C and BFP and GFP fluorescence analysed by flow cytometry (BD FACS Canto; excitation 405 nm and 488 nm; emission filter 450/50 and 530/30 nm).

## QUANTIFICATION AND STATISTICAL ANALYSIS

All biochemical assays were performed at least 3 times independently. Data were analyzed with GraphPad Prism 6.0 (GraphPad Software). Statistical significance was estimated by ANOVA or T-tests as indicated in figure legends. For Quantification ImageJ or Image Studio Lite Ver 5.2 were applied.

## DATA AND SOFTWARE AVAILABILITY

All images shown in figures 2B, 5D, S5, and S6 are available on Mendeley doi:10.17632/jbbs52642f.1

## Supplemental Information

### Equation for fitting the dissociation data of mixtures of Hsf1wt and Hsf1Δ(202-213)

When mixing Hsf1wt and Hsf1Δ(202-213), homo- and heterotrimers will form according to a binomial distribution (*na* + *mb*)^3^ with n/m being the Hsf1wt to Hsf1Δ(202-213) ratio.

Assuming that a single Hsc70 binding site is sufficient for Hsf1 dissociation and that the number of binding sites does not influence the rate of dissociation the following equation would describe the reaction:

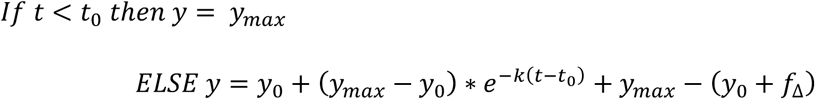

with *y_max_* and *y*_0_ representing the fitted maximal and minimal fluorescence polarization values for wild type Hsf1, *k* being the rate of the dissociation reaction, and *f*_Δ_ being the fraction of Hsf1Δ(202-213) homotrimers, which cannot be dissociated. Simulated results of this equation is shown in Fig. S4A as dashed lines.

Assuming that the number of HR-B proximal Hsc70 binding sites available in the Hsf1 trimer influences the rate by which Hsf1 timers are dissociated, the following equation system has to be used:

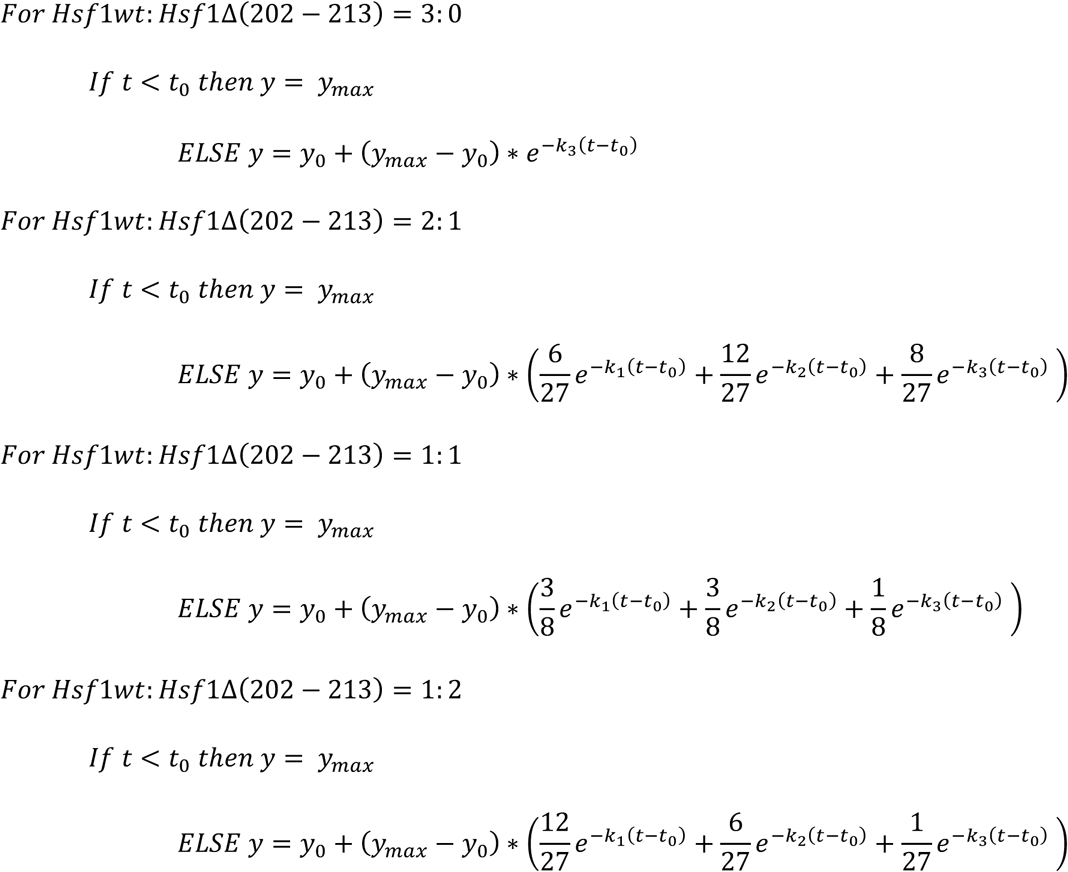

with *y_max_* and *y*_0_ representing the fitted maximal and minimal fluorescence polarization values and *k*_1_, *k*_2_, *k*_3_ being the rates of the dissociation reaction if 1, 2 or 3 Hsc70 binding sites are available per Hsf1 trimer. This equation system was fitted globally to the data of Fig. 5A. The results are shown in Fig. S4C as solid lines. Fig. S4D shows the resulting rates mean ± SD (n = 4).

## Supplemental Figures

**Figure S1 related to figure 1.**
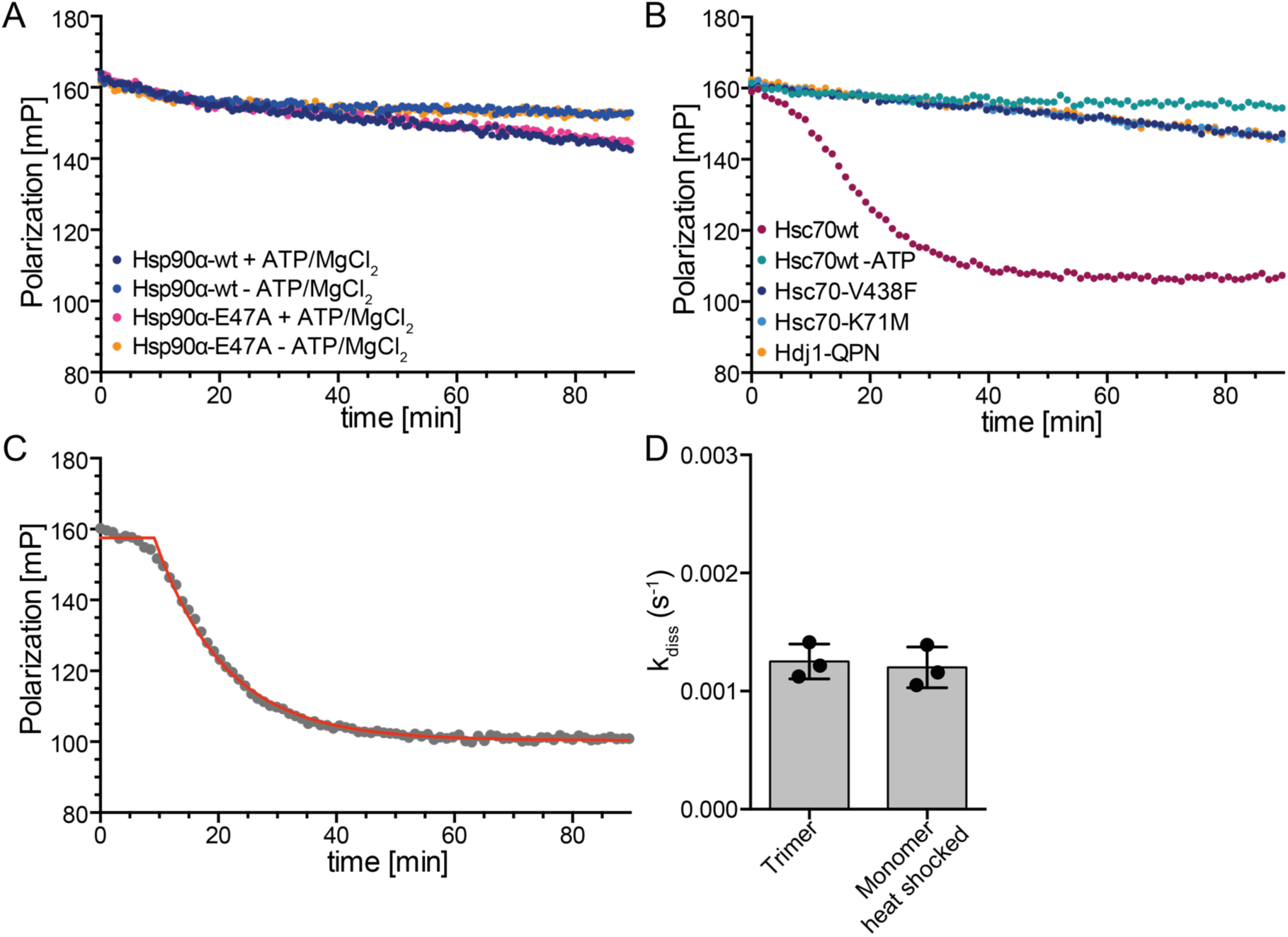
Hsc70 dissociates Hsf1 from DNA. **A**, Hsp90α wild type or ATP hydrolysis deficient Hsp90α-E47A do not influence Hsf1 DNA binding. **B**, Hsc70’s ability to hydrolyse ATP and to bind to polypeptides, and stimulation of Hsc70’s ATPase activity by DnaJB1 are essential for dissociation of DNA-bound Hsf1. Hsc70-K71M, ATPase deficient; Hsc70-V438F, reduced affinity for polypeptides; DnaJB1-H32Q,D34N (DnaJB1-QPN), defective in stimulation of Hsc70’s ATPase activity. **C**, Analysis of the dissociation rate. Fit of the composite equation [umath3] with *y_max_* and *y*_0_ representing the fitted maximal and minimal fluorescence polarization values and *k* being the rate of the dissociation reaction (red line). **D**, The dissociation rate did not differ, if Hsf1 purified as a trimer from *E. coli* without prior heat shock, or monomeric Hsf1 heat shocked for 10 min at 42°C were used for the reaction.

**Figure S2 related to figure 3.**
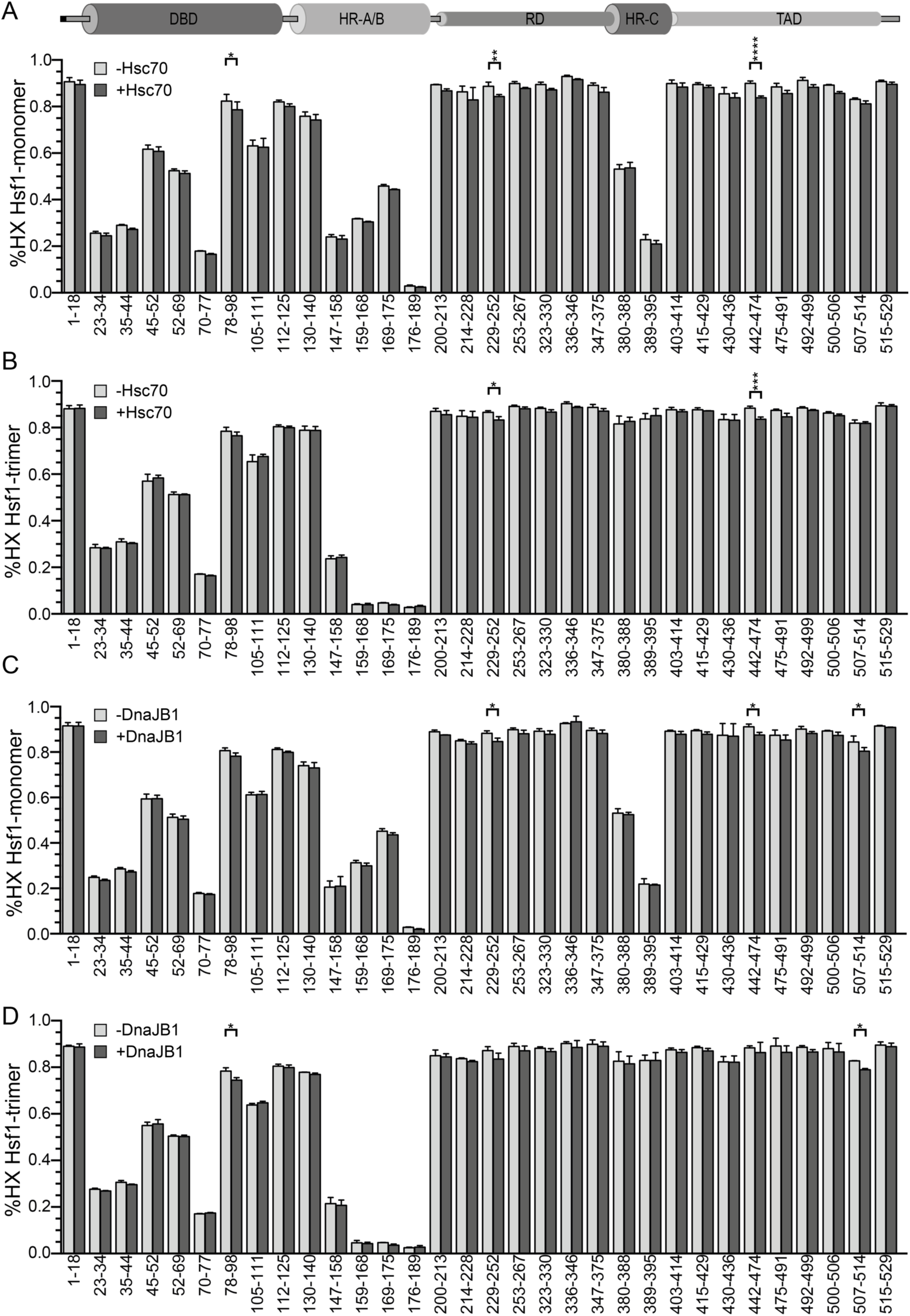
Hsc70 and DnaJB1 protect specific regions in Hsf1 monomers and trimers. Relative deuteron incorporation rates in monomeric (A, C) and trimeric (B, D) Hsf1 in the absence (light gray bars) or presence of Hsc70 (A, B; dark gray bars) or DnaJB1 (C, D; dark gray bars). Values represent means ± SD (n=3); *, p ≤ 0.05; **, p ≤ 0.01; ***, P ≤ 0.001; ANOVA, Sidak’s multiple comparison test.

**Figure S3 related to figure 3.**
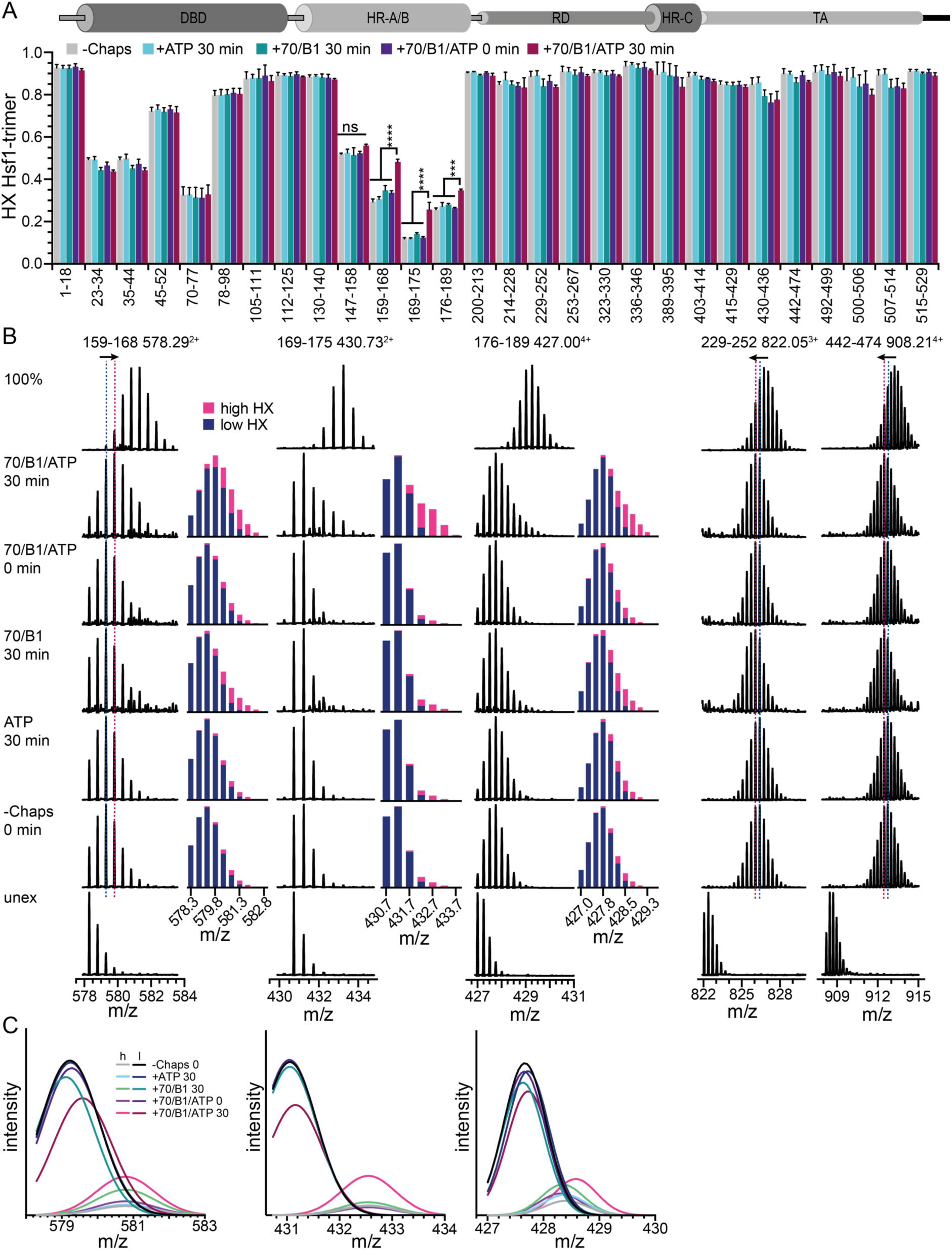
Hsc70 and DnaJB1 induce local unfolding in the trimerization domain of Hsf1. **A**, Fractional hydrogen exchange in Hsf1 segments as indicated upon incubation for 300 s at 25°C in D_2_O buffer in the absence of chaperones (-Chaps), or after a preincubation of trimeric Hsf1 in the presence of ATP, Hsc70 (70) and DnaJB1 (B1) for 0 or 30 min as indicated. The values represent mean and standard deviation (n = 3). Statistical significance was established by ANOVA with Sidak’s multiple comparison (ns, not significant; ***, p ≤ 0.001; ****, p ≤ 0.0001). On top, cartoon of Hsf1 domains to locate the exchanging segments. **B**, Original spectra of peptic peptides 578.29^3+^ (amino acids 159-168), 430.73^2+^ (aa 169-175), 427.00^4+^ (aa 176-189), 822.05^3+^ (aa 229-252), and 908.21^4+^ (aa 442-474) from unexchanged Hsf1 (unex), the 100% deuterated Hsf1, and Hsf1 incubated at 25°C for 300 s in D_2_O buffer in the absence of Hsc70, DnaJB1 and ATP (-Chaps) or after preincubation in the presence of Hsc70 (70), DnaJB1 (B1) and ATP for 0 or 30 min as indicated. The bar graphs to the right of the spectra show the fractional peak intensities of the low (dark blue) and high (magenta) exchanging subpopulation as calculated from the fit of two Gaussian distributions to the maximal isotope peak intensities (panel C and Fig. 3C). Blue and red dashed lines in spectra for peptides 578.29^3+^, 822.05^3+^, and 908.21^4+^ indicate the highest peak in the samples without chaperones and the sample with Hsc70, DnaJB1 and ATP incubated for 30 min, respectively. Arrows on top of the dashed lines indicate the Hsc70-mediated overall change in deuteron incorporation. **C**, Individual Gaussian distributions for the high (h) and low (l) exchanging species for the different conditions the sum of which results in the curves in Fig. 3C.

**Figure S4 related to figure 5.**
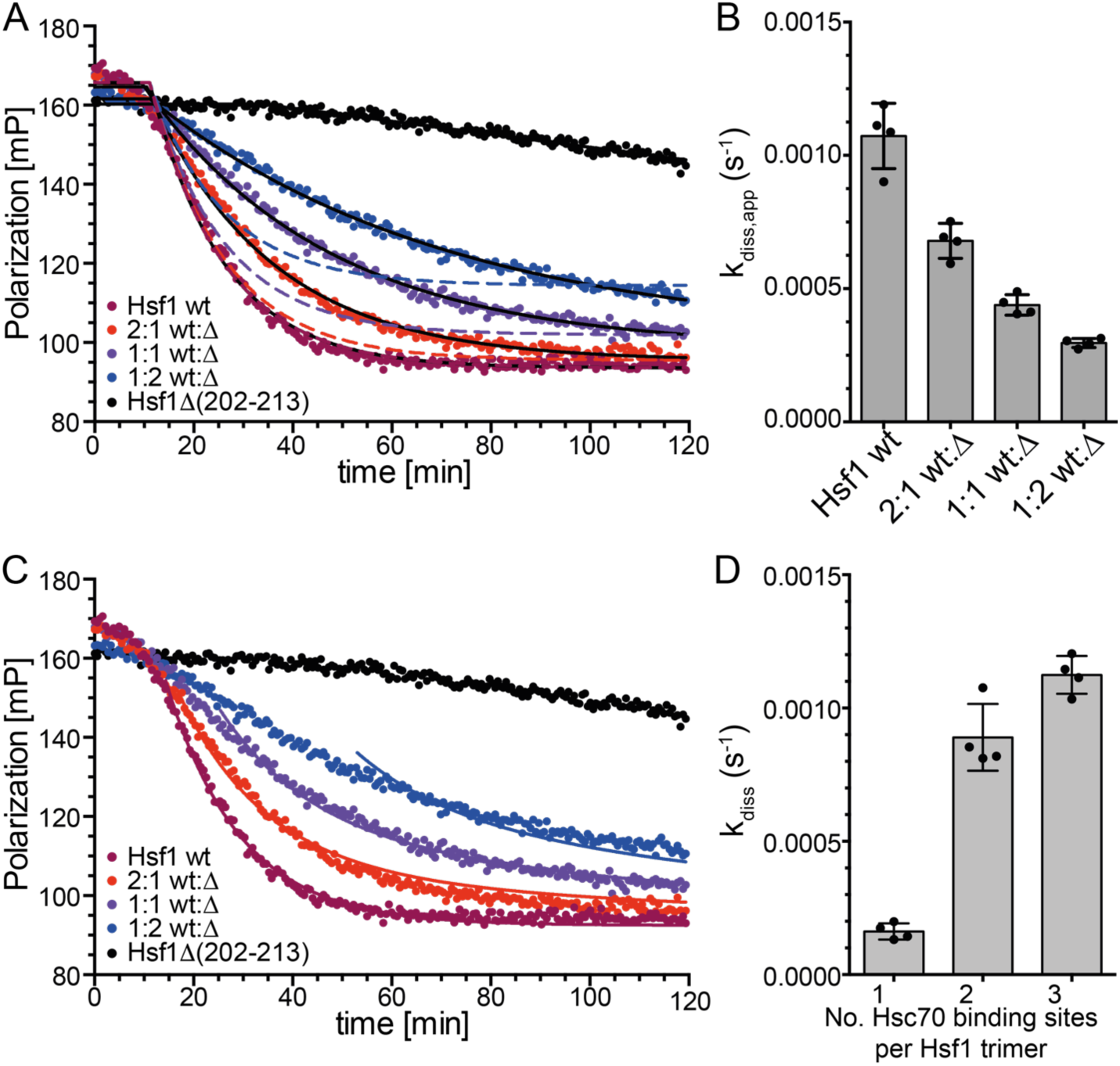
The rate of Hsc70/DnaJB1-mediated dissociation of Hsf1 from HSE-DNA depends on the number of HR-B proximal Hsc70 binding sites. **A**, Data from figure 5A fitted with the composite exponential decay function shown in fig. S1C (solid black lines). Dashed lines, theoretical dissociation functions assuming that a single Hsc70 binding site is sufficient for Hsf1 trimer dissociation. **B**, Hsf1 dissociation rates for the different mixtures fitted with function from Fig. S1C. **C**, Global fit assuming that the number of HR-B proximal Hsc70 binding sites influence the rate of dissociation of Hsf1 bound to HSE-DNA using the function shown in supplemental information. **D**, Dissociation rates derived from the global fit of panel C for Hsf1 trimers with a single (k_1_), two (k_2_), or three (k_3_) HR-B proximal Hsc70 binding sites available; mean ± SD (n=4).

**Figure S5 related to figure 5.**
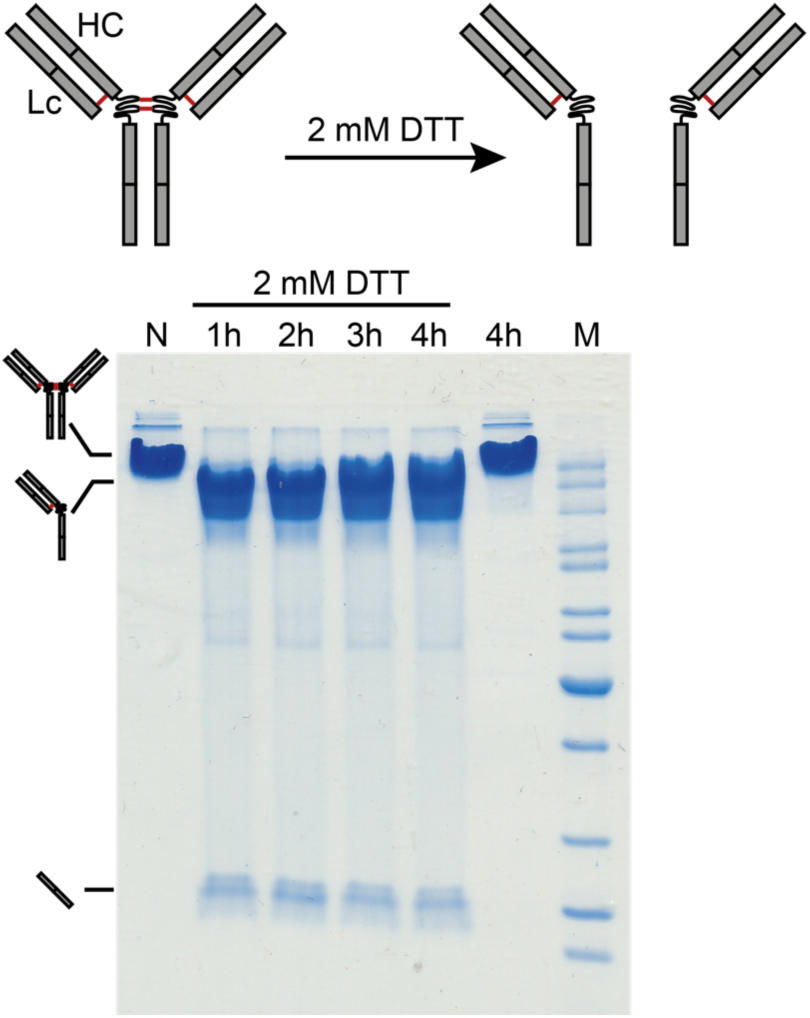
Anti-FLAG antibodies are split in halfmers by incubation with DTT. Samples were incubated in the absence (N and 4h) or in the presence of 2 mM DTT for 1 to 4 h, as indicated, separated by non-reducing SDS-polyacylamide gel electrophoresis, and the gel subsequently stained by Coomassie Brilliant Blue. M, protein size marker.

**Figure S6 related to figure 5.**
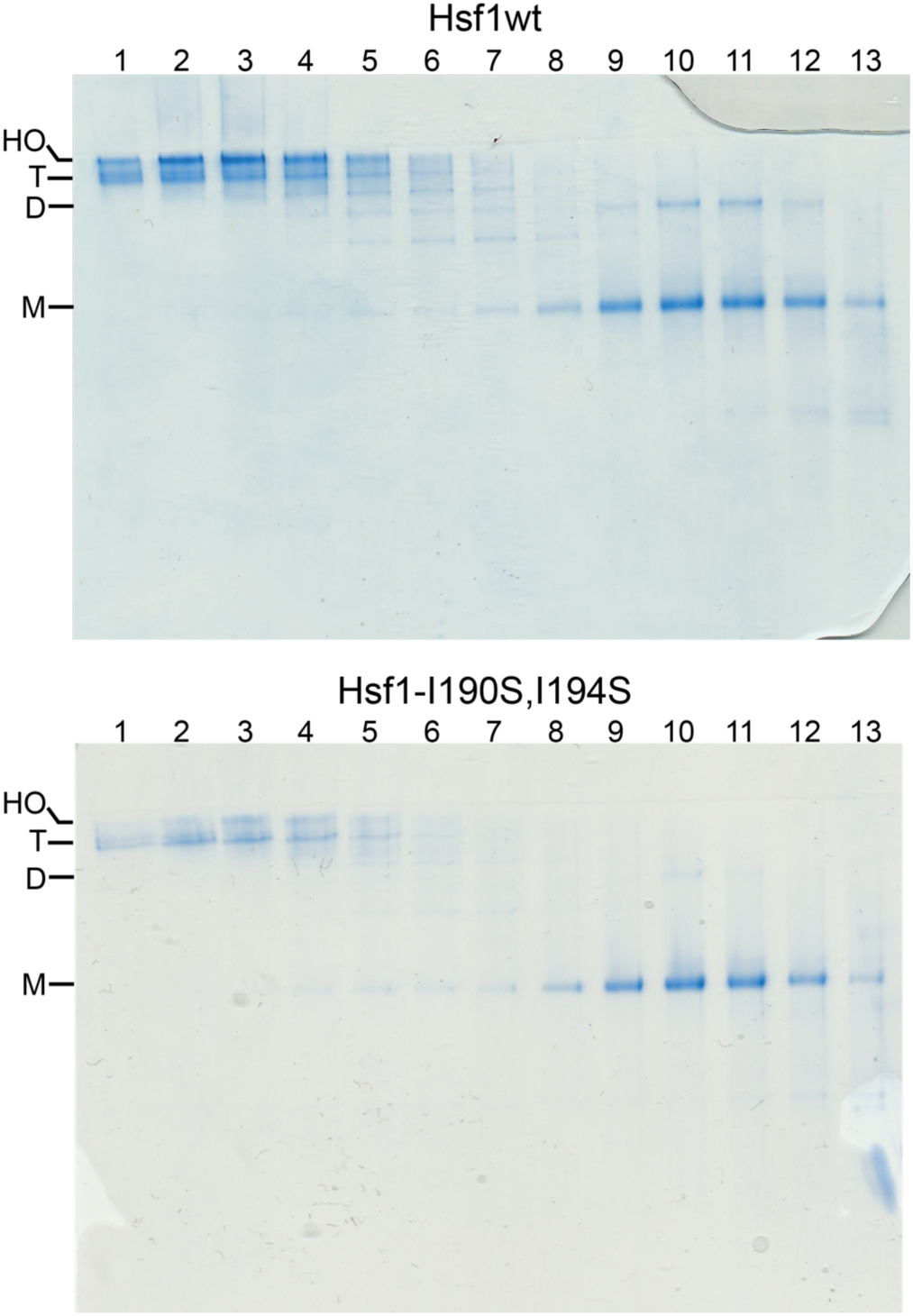
Hsf1-I190S,I194S forms less trimers and higher order oligomers than Hsf1wt. Monomeric and trimeric species of Hsf1wt (upper panel) and Hsf1-I190S,I194S (lower panel) freshly purified from an over expressing *E. coli* strain were separated by size exclusion chromatography and fractions were analyzed by BN-PAGE. Lanes 1-13 consecutive fractions from size exclusion chromatography. HO, higher order oligomers; T, Hsf1 trimers; D, Hsf1 dimers; M, Hsf1 monomers.

